## Supplementary Information for "Genetic similarity between relatives provides evidence on the presence and history of assortative mating"

#### Outline

In Supplementary Note 1, we examine the consequences of other sources of familial resemblance besides shared additive genetic effects at equilibrium. In Supplementary Note 2, we will relax the assumption of equal variance across generations, and thereby investigate the dynamics under disequilibrium. In Supplementary Note 3, we will investigate what will happen to familial similarity in polygenic indices under assortative mating, both in equilibrium and disequilibrium. In Supplementary Note 4 and 5, we provide simulations that validate our theoretical expectations. In Supplementary Note 6 and onwards, we provide other supplementary methods and information.

Because we refer to it throughout this document, we reproduce Equation (5) (aka. the algorithm) here:

$$r_{g_k} = \left(\frac{1+\rho_g}{2}\right)^k \tag{5}$$

### Supplementary Note 1 Equilibrium

This note will describe genotypic correlations at equilibrium under various assumptions. First, in Note 1.1, we introduce within-generation causes of familial similarity that comes on top of additive genetic similarity (i.e., dominant genetic effects and environmental effects shared by siblings). We show that this does not affect genetic similarity within the nuclear family but does increase genetic similarity wherever it involves two sorting processes in the same generation (e.g., first cousins). In Note 1.2, we introduce intergenerational environmental transmission in the form of cultural transmission (parental environment to offspring environment) and phenotypic transmission (parental phenotype to offspring environment). This will induce gene-environment correlations, which increase the genetic consequences of assortative mating.

##### *Key takeaways*

- At equilibrium, the relationship between the genotypic correlation between partners and the genotypic correlation between *first-degree* relatives does *not* depend on assumptions about dominance effects, environmental effects or gene-environment correlations.
- Dominance effects are unaffected by assortative mating in the polygenic model.
- Genotypic correlations between first cousins are slightly increased if there are other sources of sibling similarity in addition to shared additive genetic factors. Gene-environment correlations increase correlations between all higher-degree relatives beyond what Equation (5) in the main paper would predict. Equation (5) can still be used as a coarse approximation, although statistical tests will be biased.
- Assortative mating will induce and increase gene-environment correlations substantially if environmental transmission and genetic transmission are happening simultaneously.
- Genetic similarity between partners is greatly increased if there are gene-environment correlations (all else equal), because gene-environment correlations essentially mimic higher heritability (i.e., a larger squared correlation between genotype and phenotype). In other words, the genetic consequences of assortative mating, such as inflated genetic similarity between relatives, will be greatly exacerbated by gene-environment correlations.

##### *Some notes on what $A$ and $h$ represent*

Genetic factors are indicated by the variable $A$ in all path diagrams, which represents the sum all loci weighted by their effect on the phenotype. Summarizing all genetic loci as one variable instead of many is made possible by assuming the polygenic inheritance model[1], which is a reasonable approach to the current research problem as most complex traits seems to be highly polygenic[2]. The simulations presented in Note 5.1 shows that this approach is valid. In diagrams assuming equilibrium, the sum is rescaled to have unit variance, and $h$ represents the total standardized effect of genetic differences on phenotypic differences. Note that, because of the sum of variances law, the effect of the genetic factor will include differences caused by covariance between causal loci (i.e., linkage disequilibrium), and is as such not just the sum of direct effects. It is the induced covariance between causal loci that leads to increasing genetic variance under assortative mating (see Supplementary Note 2).

#### 1.1 Dominance effects and environmental effects shared by siblings

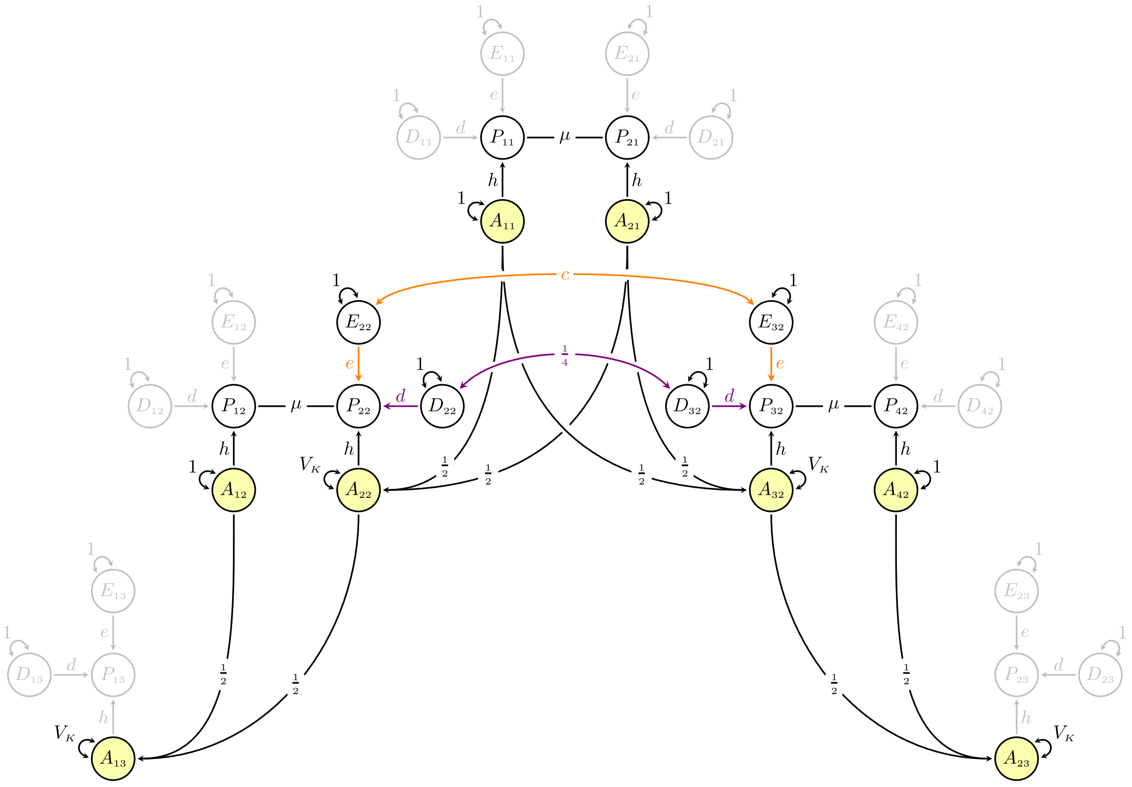

**Supplementary Figure 1** *This path diagram adapts Fig. 1 by adding other causes of sibling similarity, namely dominance effects (purple) and sibling-correlated environmental effects (orange). Pathways that are irrelevant for understanding genetic similarity among the depicted individuals are greyed out to remove clutter.*

In Supplementary Figure 1, we have adapted Fig. 1 from the main paper by adding a correlation (denoted $c$, coloured orange) between siblings' environmental effects. The shared environmental influences are generation-specific and is therefore independent of the parental phenotype or parental environment (Intergenerational environmental transmission are discussed in Note 1.2). In classical twin designs, environments shared by siblings are usually parameterized as its own variable that is perfectly correlated between siblings[3]. However, to keep the model simple and more generalizable, we have instead denoted it with a correlation, $c$, between the existing environmental variable ($E$). The phenotypic correlation between siblings that are attributable to shared environments will here be $ce^2$, meaning $c$ can also be read as the proportion of environmental variance that is shared between siblings.

The diagram in Supplementary Figure 1 also includes dominance deviation effects, denoted $D$ with effect $d$ (coloured purple) on the phenotype. Dominance effects are non-additive genetic effects where the difference between having 0 and 1 copies of an allele does not equal the difference between having 1 and 2 copies of an allele[3]. It is characterized by the average phenotype of heterozygotes (i.e., 1 copy) deviating from the midpoint between the two homozygotes (0 and 2 copies). If the trait in question is influenced by more than just a few genetic loci, then dominance variance and the resulting familial similarity will be practically unaffected by assortative mating[1,4]. The dominance correlation between parent and offspring will be 0 and the dominance correlation between siblings will be $\frac{1}{4}$. This is reflected in how the path diagram in Supplementary Figure 1 is drawn.

##### *Multiple assortment processes open new pathways*

If we trace all valid paths between partners and between first-degree relatives using the diagram in Supplementary Figure 1, we will end up with the same equations as before. The same goes for the avuncular genotypic correlation and the grandparent-grandchild genotypic correlation. There are no new pathways between the additive genetic components, meaning dominance effects and sibling-shared environmental effects have no consequence on the additive genetic similarity between these family members. However, there are new pathways between partners-of-siblings-in-law (henceforth, *co-siblings-in-law*), and therefore new pathways between first cousins. These new pathways have in common that they are taking two assortment processes into account in the same generations (e.g., mother–father and aunt–uncle). Because of the special path tracing rules pertaining to copaths, where the traditional path tracing rules are effectively reset whenever the co-path is traversed, it is possible to have valid pathways between the two sorting processes that completely bypass additive genetic influences. The genetic factors of co-siblings-in-law ($A_{12}$ and $A_{42}$) can therefore be correlated via pathways that go through dominance covariance (purple) or environmental covariance (orange) between siblings. This is not accounted for in the algorithm described in the main paper (i.e., $r_{g_{co-in-laws}} = r_{g_1}{\rho_g}^2$), which would therefore underestimate the true genotypic correlation.

##### *Finding the new correlations between co-siblings-in-law and first cousins*

Rather than explicitly defining all the ways in which siblings can be similar for reasons other than additive genetic factors, which in Supplementary Figure 1 would be $ce^2 + \frac{d^2}{4}$, we can define this component through exclusion. That way, we can remain agnostic as to what these influences are. If we let $r_{p_s}$ be the phenotypic correlation between siblings, then the part of the correlation that is *not* due to additive genetic factors is $r_{p_s} - h^2 r_{g_1}$. The genotypic correlation between co-siblings-in-law can now be described by two components: One that is mediated by additive genetic factors shared by siblings, and one that results from other causes of similarity between siblings:

$$r_{g_{co-in-laws}} = r_{g_1}{\rho_g}^2 + \left(r_{p_s} - h^2 r_{g_1}\right)\mu\rho_g \qquad \text{(S1.1)}$$

This second component will also cause resemblance in the following generations, meaning the genotypic correlation between first cousins in the presence of such extra pathways is:

$$r_{g_{cousins}} = r_{g_1}^3 + \frac{\left(r_{p_s} - h^2 r_{g_1}\right)\mu\rho_g}{4} \qquad \text{(S1.2)}$$

##### *Some notes on dominance effects under assortative mating.*

While Fisher [1] did not consider cases where siblings resemble each other due to correlated environments, he did consider cases where siblings resemble each other due to effects of dominance. If we consider a case where siblings are correlated due to additive genetic effects and dominant genetic effects ($r_{p_s} = h^2 r_{g_1} + \frac{d^2}{4}$), the genotypic correlation between first cousins becomes:

$$r_{g_1}^3 + \frac{d^2\mu\rho_g}{16} \qquad \text{(S1.3)}$$

This can be verified by re-tracing the genotypic correlation between cousins in Supplementary Figure 1 while assuming $c = 0$. If we multiply Equation (S1.3) with the heritability, it equals the expected phenotypic correlation as reported in Fisher [1] and later Lynch and Walsh [4]. However, as we can see here, dominance covariance is just a special case of resemblance between siblings that is not attributable to shared additive genetic factors and is included in $r_{p_s} - h^2 r_{g_1}$. It is not the dominance effects that become correlated, but the additive genetic effects that become further correlated. This is not clear in the original sources, where readers can be forgiven for thinking that it is the dominance components of cousins that become correlated under assortative mating. Using path diagrams to understand these correlations makes it obvious that this is not the case.

##### *What would be the consequences?*

Dominance variance is rarely a substantial contributor to phenotypic variance[5], and because siblings only share $\frac{1}{4}$ of their dominance variance, the increased genotypic correlation between cousins owing to dominance variance will in most circumstances be negligible (we confirmed this with simulations, see Note 5.2). The same goes, albeit to a lesser extent, for the environmental pathways. For example, for a trait where $h^2 = 40\%$, $\mu = .50$, and $c = \frac{2}{3}$ (meaning $40\%$ of the variance is caused by shared environmental influences), the genotypic correlation between cousins should be $r_{g_{cousins}} = .226$, but Equation (5) would predict $r_{g_1}^3 = .216$ (we confirmed this with simulations, see Note 5.3). This difference of .010 (4.6%) is detectable with enough statistical power and may bias a statistical test that relies on Equation (5) (e.g., testing equilibrium, see Supplementary Note 2), but Equation (5) still serves as a reasonably good approximation. For co-siblings-in-law, one the other hand, the difference will be larger. In the above example, the genotypic correlation will be $r_{g_{co-in-laws}} = .064$, which is more than twice what it would have been without the shared environmental effects.

##### *Half-siblings are also affected by multiple assortment processes*

Even though half-siblings are second-degree relatives, it is not possible to use Equation (5) to obtain the expected genotypic correlation. Two sorting processes are involved, which leads to the same problems as we encountered when siblings where similar for non-genetic reasons. Supplementary Figure 2 shows a path model where one person in the parent generation ($P_{21}$) had offspring with two different partners ($P_{11}$ and $P_{31}$, respectively, which we will call the stepparents). This means meaning that $P_{12}$ and $P_{22}$ are half-siblings. If we trace the pathways between the two stepparents' genetic factors ($A_{11} \leftrightarrow A_{31}$), we find two valid chains: (1) $h\mu hh\mu h$ and (2) $h\mu ee\mu h$. The first chain is linked through the genetic component of the shared parent's phenotype, whereas the second component is linked through the non-genetic component. After simplification, we can write:

$$r_{g_{step-parents}} = {\rho_g}^2 + \mu\rho_g e^2 \tag{S1.4}$$

This second component will also cause resemblance in following generations, meaning the genotypic correlation between half-siblings will be:

$$r_{g_{half-siblings}} = r_{g_1}^2 + \frac{\mu\rho_g e^2}{4} \tag{S1.5}$$

Unlike most other relatives, the expected genotypic correlation between half-siblings cannot be stated in terms of the genotypic correlation between partners alone but requires the phenotypic correlation as well. We can simplify (S1.5) to $\frac{1+2\rho_g+\mu\rho_g}{4}$, which, when multiplied by the heritability, equals the phenotypic correlation reported by Nagylaki [6] and later Lynch and Walsh [4]. Note also that half-siblings may lead to other complications which should be accounted for. For example, the strength of assortment may be different for first and second partnerships, and the phenotype that is sorted upon may change across time [7]. We urge researchers to draw the path diagram that would be relevant for their own research and re-derive the relevant equations. For this paper, we conclude that half-siblings bring with them too many complications as to be informative for the current research questions.

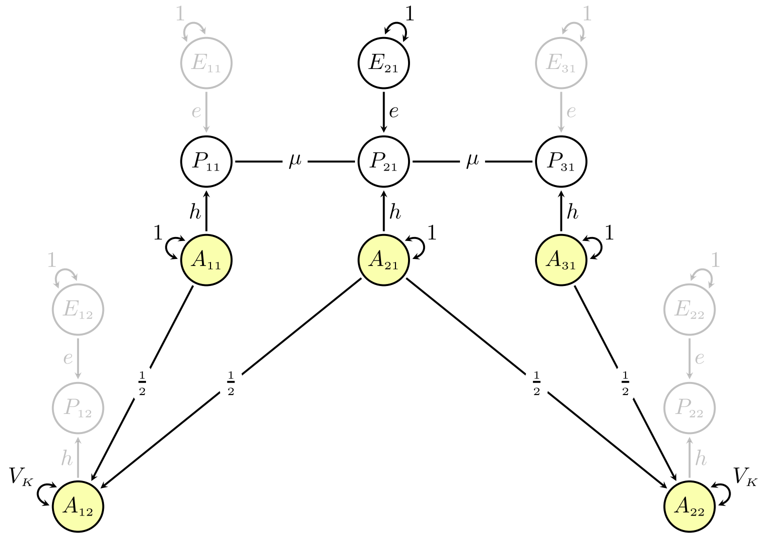

**Supplementary Figure 2** *Path diagram of genetic similarity between half-siblings under assortative mating (assuming genetic transmission only). Pathways that are irrelevant for understanding genetic similarity among the depicted individuals are greyed out to remove clutter.*

#### 1.2 Environmental transmission and gene-environment correlations

Up to now, we have only considered *genetic transmission* across generations. In this section, we consider two forms of environmental transmission as well. First, we will consider what we call *cultural transmission*, which is when the parental environment forms part of the offspring environment (e.g., $E_{11} \rightarrow E_{22}$)[8]. We will then also add *phenotypic transmission*, which is when the parental phenotype forms part of the offspring environment (e.g., $P_{11} \rightarrow E_{22}$). We will then examine how this affects genetic similarity between various family members.

##### *Cultural transmission*

Supplementary Figure 3 extends Fig. 1 by adding causal effects (denoted $b$, coloured red) of the parental environment on the offspring environment. Note that each parent has separate causal effects, meaning the combined causal effect of both parents' environments would be $2b$ (in addition to any covariance between the two). This is completely analogous to genetic transmission (coloured blue), the only difference being that the genetic equivalent of $b$ is known to be exactly one half[1]. As noted by Rice, Cloninger and Reich, if $b$ happens to be close to one half, then genetic and cultural transmission will be indistinguishable in extended families[9].

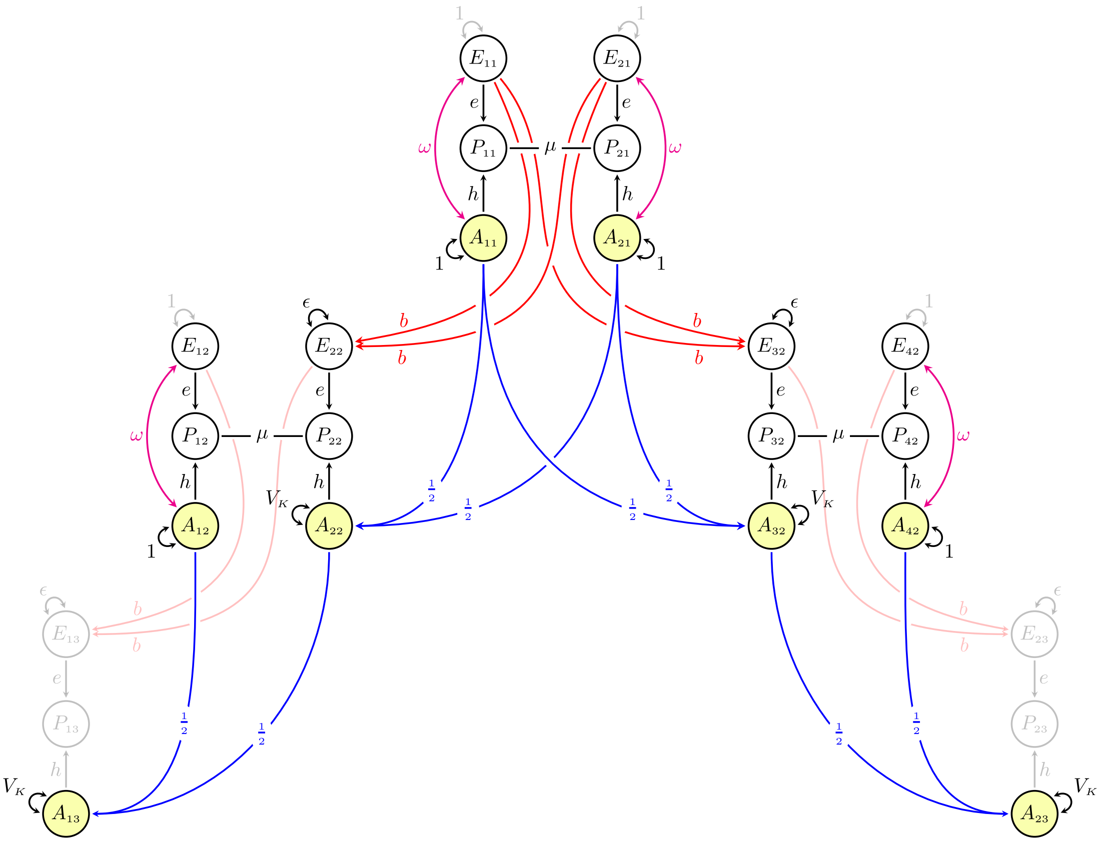

**Supplementary Figure 3** *This path diagram adapts Fig. 1 by including both genetic (blue) and cultural (red) transmission across generations. Assortative mating will here induce correlations between genetic and environmental influences (magenta), which we for simplicity's sake assume to be constant across generations. Pathways that are irrelevant for understanding genetic similarity among the depicted individuals are greyed out to remove clutter.*

In the absence of assortative mating, genetic and cultural transmission can occur independently of each other. However, under assortative mating, genetic influences inherited from the mother will become correlated with the environmental influences transmitted from the father (and vice versa). In other words, assortative mating will induce a gene-environment correlation. The exact magnitude of this correlation will depend on the gene-environment correlation in the preceding generations (meaning gene-environment correlations, like assortative mating, will increase phenotypic variance and phenotypic resemblance until it too reaches an equilibrium, see Note 2.2). For simplicity, we will for now assume that the gene-environment correlation is also at equilibrium (i.e., it is stable across generations), although we acknowledge that this may be a less reasonable assumption in many circumstances than for assortative mating. This assumption is instantiated in two ways: First, all variables have unit variance (meaning variance is constant across generations). Second, the gene environment correlation is defined to be equal across generations. For individuals whose parents are not drawn in the diagram, we have added a correlation denoted $\omega$ (coloured magenta) between their genetic and environmental influences. To find the gene-environment correlation, we can trace all valid pathways between the genetic and environmental influences of an individual whose parents are included in the diagram (e.g., $A_{22} \leftrightarrow E_{22}$). After simplification, we are left with:

$$\omega = \omega b + (h + \omega e)\mu(e + \omega h)b \qquad \text{(S1.6)}$$

As we can see, the gene-environment correlation consists of two components, one that depends on the gene environment correlation in the preceding correlation, and another that depends on assortative mating in the preceding generation. Conveniently, the gene-environment correlation happens to also equal the correlation between one individuals' genotype and their sibling's environment, as well as the correlation between an offspring's environment and their parent's genotype. (You can double-check this by, for example, tracing $E_{22} \leftrightarrow A_{11}$ or $A_{22} \leftrightarrow E_{32}$). We have used this fact to simplify some of the equations reported later.

##### *Phenotypic transmission*

Supplementary Figure 4 extends Supplementary Figure 3 by adding causal effects (denoted $p$, coloured teal) from parental phenotypes to offspring environments. Just like with cultural transmission, each parent has separate causal effects, meaning the combined causal effect of both parents would be $2p$ (in addition to any covariance between the two). Note also that the causal effect of a parental phenotype on the offspring phenotype becomes $pe$, as the effect must be weighted by the strength of environmental influences on the offspring phenotype.

Unlike cultural transmission, phenotypic transmission will induce gene-environment correlations regardless of whether assortative mating occurs or not. This is because the parental phenotype is itself partly caused by genetic variants which the offspring also inherits. For this reason, phenotypic transmission is sometimes called *genetic nurture* or *indirect genetic effects*[10-12], although we prefer the more general term "phenotypic transmission" as it does not limit its focus to the heritable component of the parent's phenotype. To find the new gene-environment correlation, we trace all pathways between, say, $A_{22}$ and $E_{22}$, and simplify:

$$\omega = \omega b + (h + \omega e)\mu(e + \omega h)b + (h + \omega e)(p + p\mu) \qquad \text{(S1.7)}$$

The first two (red) components are the same as in Equation (S1.6) (i.e., caused by cultural transmission), whereas the last (teal) component is induced by phenotypic transmission. From this equation, we can see that the gene-environment correlation is a function of assortative mating, and that stronger assortative mating means larger gene-environment correlations. If we for a moment focus exclusively on the component induced by phenotypic transmission (i.e., $b = 0$) and factor out the strength of assortment, $(hp + \omega ep)(1 + \mu)$, we see that the gene-environment correlation is inflated by a factor of $(1 + \mu)$ in the presence of assortative mating. For example, if $\mu = .50$, then $\omega$ is 50% larger than it would have been otherwise, all else equal. Note that this is at equilibrium, and that the other terms in the equation, such as $h$ and $\omega$, will also change with changing strength of assortment (as will the variances, which must then be included in the above equation).

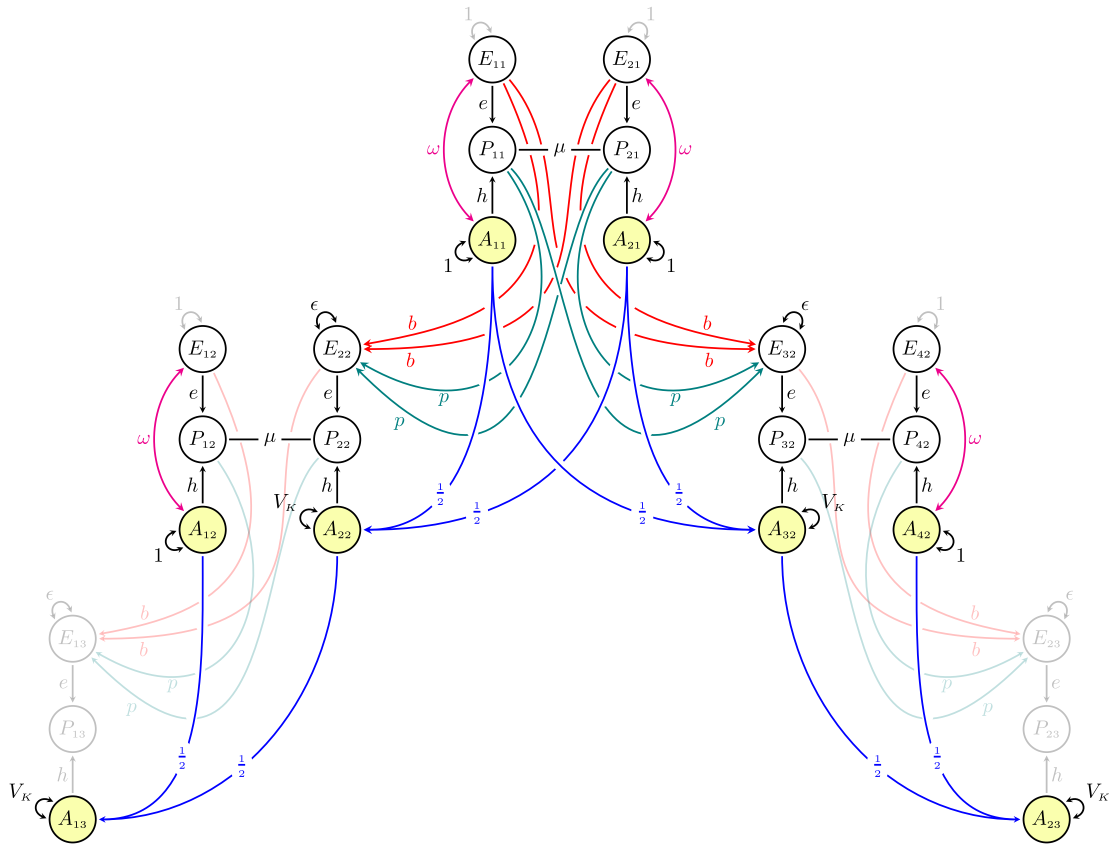

**Supplementary Figure 4** *This path diagram adapts Supplementary Figure 3 by also adding phenotypic (teal) transmission, in addition to genetic (blue) and cultural (red) transmission. Phenotypic transmission and assortative mating will here induce correlations between genetic and environmental influences (magenta), which we for simplicity's sake assume is constant across generations. Pathways that are irrelevant for understanding genetic similarity among the depicted individuals are greyed out to remove clutter.*

##### *Genotypic correlations in the nuclear family in the presence of gene-environment correlations*

We will use a prime (′) to denote genotypic correlations where gene-environment correlations have been incorporated. The genotypic correlation between partners will be different when there are gene-environment correlations because the father's genetic factors will be correlated with the mother's environment, which in turn is correlated with the mothers' genetic factors (and vice versa). If we trace the genotypic correlation between partners (e.g., $A_{11} \leftrightarrow A_{21}$) in Supplementary Figure 4, we find that:

$$\rho_g' = \mu(h + \omega e)^2 \tag{S1.8}$$

In other words, gene-environment correlations will mimic the effect of higher heritability for the genotypic partner correlation. Note also that, if the genetic effects where to suddenly disappear (i.e., $h = 0$), a gene-environment correlation would still result in a true genotypic correlation between partners, which in turn would have the same genetic consequences as if there was true genetic effects. (The gene-environment correlation would likely not be stable across generations in this scenario, see Supplementary Note 2, although it illustrates a point: It is the genotype-phenotype correlation that matters, not the heritability per se).

The genotypic correlation between first-degree relatives will be similarly altered, but if defined in relation to the genotypic correlation between partners, the equation remains the same as before. If we use path tracing on Supplementary Figure 4, we find it is still proportional to the genotypic correlation between partners:

$$r'_{g_1} = \frac{1+\rho'_g}{2} \tag{S1.9}$$

As is apparent in Equation (S1.8) and (S1.9), the genotypic correlation between partners (and consequently first-degree relatives) becomes substantially higher if genetic influences are correlated with environmental influences. For example, if $\mu = .50$, $h^2 = .40$, $b = .20$, and $p = .10$, which in turn would mean that $e^2 = 33.9\%$ and $\omega = .35$, then the genotypic correlation between partners would be $\rho_g{}' = .352$. This is 76% higher than it would have been in the absence of gene-environment correlations ($\mu h^2 = .20$). Similarly, the genotypic correlation between first-degree relatives would be $r'_{g_1} = .676$ instead of $.60$. Note that the gene-environment correlation and its consequences are relatively large even though the degree of environmental transmission is modest. For these example values, the direct causal effect of a parental phenotype on the offspring phenotype is only $pe = .058$, and a modest 7.8% of the variance can be ascribed to intergenerationally shared environmental factors[a]. In other words, even modest environmental effects may lead to considerable gene-environment correlations, and consequently large increases in genotypic correlations between relatives.

Importantly, because the genotypic correlation between first-degree relatives is proportional to the genotypic correlation between partners regardless of gene-environment correlation, it is possible to use the observed genotypic correlation between partners and first-degree relatives to test for equilibrium while remaining agnostic about the presence of gene-environment correlations. This, as we shall see, is not the case for more distant relatives.

##### ***Complications for the extended family in the presence of gene-environment correlations***

The genotypic correlations between higher-degree relatives will, like first-degree relatives, be substantially inflated in the presence of gene-environment correlations. However, the simple algorithm we established in Equation (5) will no longer be accurate. Here is why: First, gene-environment correlations open pathways between the genetic factors of higher-degree relatives that are not mediated by the genetic factors of intermediate first-degree relatives. For example, a child will inherit genes from their mother which are correlated with their father's familial environment through assortment, which in turn is correlated with their uncle's genetic factors. Environmental transmission also makes siblings similar for environmental reasons, which open pathways between two assortment processes that completely bypass genetic factors (as we saw in Note 1.1). Second, some of the pathways that connect partners' genetic factors cannot be continued without breaking path tracing rules, and thus cannot be part of a longer chain. If we mindlessly use Equation (5), we will both include invalid paths and exclude valid paths. Usually, the result is an underestimate of the true genotypic correlation. In short, neither of the properties that allowed the use of the general algorithm (Equation (5)) are still present, and we must instead retrace the paths manually.

Gene-environment correlations under assortative mating quickly leads to long and complicated equations for the genotypic correlations between higher-degree relatives. To keep the equations at a manageable length, it is useful to define a few shortcuts which we can use when we simplify the equations. We will also use different colours for the different terms to improve clarity. We have already defined the gene-environment correlation, $\omega$, which we know also equals the correlation between one individuals' genotype and their sibling's environment, as well as the correlation between an offspring's environment and their parent's genotype. Another set of pathways that reoccurs often is $(h + \omega e)\mu$, which

[a] See $r_{e_s}$ (Equation (S1.12)) below. To get variance explained, multiply by squared environmental effect: $r_{e_s}e^2$

is the correlation between an individual's genotype (A) and their partner's phenotype (P). We will denote this $\rho'_{gp}$, so that:

$$\rho'_{gp} = Corr(A_{12}, P_{22}) = (h + \omega e)\mu \tag{S1.10}$$

Using this, we can also define the part of the genotypic parent-offspring correlation that *can* form part of a longer pathway. This is the set of pathways that end at the parent's genotype while still tracing backwards against the direction of the arrows. We will denote this $r_{g_\psi}$ (for no better reason that that $\psi$ has a line that continues above and beyond, symbolizing that it can form part of longer pathways):

$$r'_{g_\psi} = \frac{1 + h\rho'_{gp}}{2} \tag{S1.11}$$

Finally, we will define $r_{e_s}$ as the correlation between sibling environmental factors (i.e., $E_{22} \leftrightarrow E_{32}$). In the path diagram in Supplementary Figure 4, we find that this is attributable to three sets of pathways: (1) that attributable to cultural transmission, (2) that attributable to phenotypic transmission, and (3) that attributable to the correlation between cultural and phenotypic transmission:

$$r_{e_s} = 2b^2(1 + \mu(e + \omega h)^2) + 2p^2(1 + \mu) + 4bp(e + \omega h)(1 + \mu) \tag{S1.12}$$

##### ***Genotypic correlations in the extended family under cultural transmission***

Following Supplementary Figure 3, we can trace the avuncular genotypic correlation (e.g., $A_{13} \leftrightarrow A_{32}$), which ends up simplifying to:

$$r'_{g_2} = r'_{g_\psi} r'_{g_1} + \frac{e\omega\rho'_{gp}}{2} \tag{S1.13}$$

The first component takes care of the invalid pathways, and the second part takes care of pathways that bypass the genetic factors of first-degree relatives. The genotypic correlation between grandparents and offspring (e.g., $A_{11} \leftrightarrow A_{13}$) is the same as Equation (S1.13). For first cousins, the correct equation after simplification becomes:

$$r'_{g_{cousins}} = r'_{g_1} {r'_{g_\psi}}^2 + \frac{2r'_{g_\psi}\omega e\rho'_{gp}}{2} + \frac{e^2 r_{e_s} {\rho'_{gp}}^2}{4} \tag{S1.14}$$

The equation has three components: (1) pathways that are mediated by genetic similarity between first-degree relatives, (2) additional pathways that are mediated by genetic similarity between second-degree relatives, and (3) pathways that completely bypass genetic similarity of both first- and second-degree relatives. We leave it as an exercise to the reader to trace the genotypic correlation between great-grandparents (not drawn) and offspring, although note that it does not equal the genotypic correlation between first cousins despite both being third-degree relatives.

As is apparent, it is not possible to state the expected genotypic correlation for higher-degree relatives in simple equations, nor in terms of the genotypic correlation between partners alone. Despite these complications, using Equation (5) is still a reasonably good approximation. In the above example ($\mu = .50$, $h^2 = .40$, $b = .20$ and $p = .10$), the true genotypic correlation between first cousins will be $r'_{g_{cousins}} = .334$, whereas it would be approximated to $.309$ if one used Equation (5). Despite being a decent approximation, the inaccuracy is enough to cause bias in statistical tests that assume $r_{g_{cousins}} = r^3_{g_1}$, such as testing equilibrium.

In Note 5.4, we provide simulations that verify the theoretical expectations laid out here.

#### Supplementary Note 2 Disequilibrium

This note will describe genotypic correlations under disequilibrium. To do this, we must first describe changes in genetic variance across generations. In Note 2.1, we focus on a model with genetic transmission only, showing that genetic variance increases towards an equilibrium as a function of the genotypic correlation between partners in the preceding generation. In Note 2.2, we include environmental transmission and describe how both the genetic variance, environmental variance, and gene-environment correlation will increase towards an equilibrium with successive generations of assortative mating. Finally, in Note 2.3, we consider the correlation between parents and offspring during disequilibrium. There are several reasons for this only focusing on the parent-offspring correlation: First, as we have seen, the genotypic correlations between distant relatives will depend on the presence of environmental effects and, for polygenic indices, the genetic signal. Using them to test for equilibrium – which is the aim of this paper – therefore requires strong assumptions about genetic signal and degree of gene-environment correlation which we are not willing to make. Second, we have many more parent-offspring dyads (117,041) in our data than other dyads (e.g., siblings: 22,575), meaning this correlation will be estimated with much higher precision and consequently provide the most high-powered test of equilibrium. Third, focusing on parent-offspring correlations rather than sibling correlations removes the need to distinguish between siblings in the parent generation and siblings in offspring generation. This would be required when investigating disequilibrium using siblings, which in turn would reduce the effective sample size.

##### *Key takeaways*

- Genetic variance in generation $t + 1$ depends on (1) the genetic variance in generation $t$, (2) the genotypic correlation between partners in generation $t$, and (3) the recombination variance.
- Genetic variance will increase asymptotically towards an equilibrium, which is reached after 6 to 10 generations. Higher heritabilities or stronger partner correlations leads to more generations in disequilibrium.
- Gene-environment correlations mimic higher heritabilities, leading to a stronger genotypic correlation between partners (all else equal), which in turn leads to a larger increase in genetic variance and more generations in disequilibrium.
- Heritability usually increases as well, although if there is environmental transmission as well, environmental variance and variance attributable to gene-environment covariance may increase more quickly, which would lead to decreasing heritability.
- The genotypic correlations between relatives such as parents and offspring also increase toward an equilibrium.
- During disequilibrium, Equation (5) overestimates the true correlation between relatives. A mismatch between the true correlation and what Equation (5) would predict is therefore evidence of disequilibrium.
- The degree of mismatch depends on the number of generations of assortative mating.

#### 2.1 Genetic variance across generations

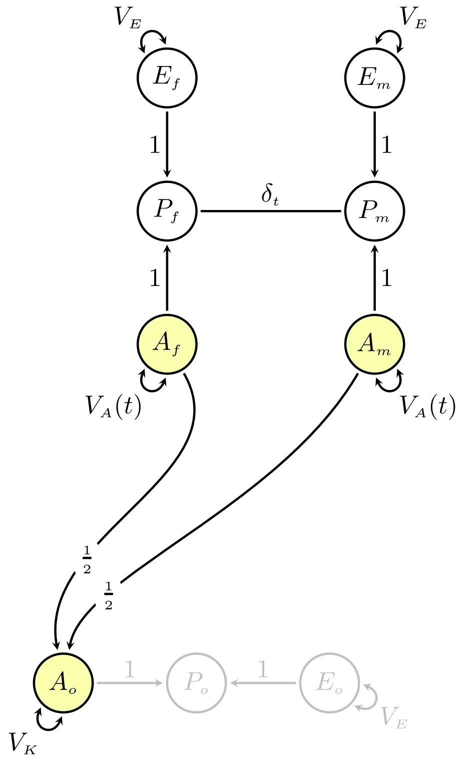

**Supplementary Figure 5** *Unstandardized path diagram of genetic inheritance under assortative mating.*

Because assortative mating will change genetic variance before equilibrium, we can no longer assume unit variances as we did in the previous note. Supplementary Figure 5 therefore uses explicit variance components. This changes how we derive correlations, because tracing all valid chains between two variables will give us the covariance, not the correlation. To turn a covariance into a correlation, we need to divide by the geometric mean of the two variables' variances. To find the variances, we must trace all valid chains leading from the two variables back to themselves.

$$\frac{Cov(X,Y)}{\sqrt{Var(X) \times Var(Y)}} \tag{S2.1}$$

To understand the correlation between relatives under disequilibrium, we must therefore first understand how the variance changes. The equations for changes in genetic variance across generations are available elsewhere[13-15], but because we will be extending the model to see how gene-environment correlations and the use of polygenic indices affects this process, it will be useful to re-derive them from first principles using path analysis.

We let $t$ denote the number of *prior* generations of assortative mating, where $t = 0$ is the generation where partners first match on the phenotype. Note that this would imply that partners in generation $t = 1$ (the first generation *after* assortment began) would be the *second* generation to assort on the phenotype. Supplementary Figure 5 shows a mother-father-offspring trio (subscripted $m$, $f$, and $o$, respectively) where the parents are in generation $t$. The genetic variance in generation $t$ is denoted $V_A(t)$, and the recombination variance is denoted $V_K$. The recombination variance will depend on the genetic variance in the base population ($t = 0$, before assortative mating) and will be[13-16]:

$$V_K = \frac{V_A(0)}{2} \tag{S2.2}$$

The genetic and environmental *effects* are for simplicity's sake assumed to be constant across generations and therefore set to 1. We are also assuming that the environmental variance, denoted $V_E$, is constant across generations, and that there are no sex differences. If you want to relax these assumptions, you can replace them by appropriate parameters (e.g., $V_E(t)$) and redo the path tracing with these changes.

As always, the heritability in generation $t$ is the proportion of phenotypic variance attributable to genetic variance:

$$h^2(t) = \frac{V_A(t)}{V_A(t) + V_E} \quad (S2.3)$$

The partner similarity is slightly more complicated because the copath coefficient is neither the covariance nor the correlation between partners when variances are not equal to one. Instead, it is a special coefficient denoting the strength of assortment. Because we have already defined $\mu$ as the phenotypic *correlation*, the assortment strength in generation $t$ is now denoted $\delta_t$ to avoid confusion. We can use path tracing to find the phenotypic partner correlation:

$$\mu = \frac{\delta_t(V_A(t) + V_E)^2}{V_A(t) + V_E} = \delta_t(V_A(t) + V_E) \quad (S2.4)$$

We can apply similar logic to the genotypic correlation between partners:

$$\rho_g(t) = \frac{\delta_t V_A^2(t)}{V_A(t)} = \delta_t V_A(t) \quad (S2.5)$$

Because heritability will change across generations, all else being equal, so will the genotypic correlation between partners – even when keeping the phenotypic correlation constant. Note also that we can transform Equation (S2.5) using Equation (S2.3) and Equation (S2.4) and find that $\rho_g(t) = \mu h^2(t)$, thus matching Equation (1) in the main paper. Finally, to keep $\mu$ constant while $V_A$ increases, $\delta_t$ will have to decrease accordingly. It is an open question whether it is more realistic to keep $\mu$ or $\delta_t$ constant when investigating assortative mating across generations, but here we follow previous work and let $\mu$ be constant[6,13]. This does not affect the equations, but it would change the resulting figures somewhat.

We are now ready to derive the genetic variance in the offspring generation. We do that by tracing all paths leading from $A_o$ back to itself:

$$V_A(t+1) = Var(A_o)$$

$$= V_K + \frac{V_A(t)}{4} + \frac{V_A(t)}{4} + \frac{\delta_t V_A^2(t)}{4} + \frac{\delta_t V_A^2(t)}{4}$$

$$= V_K + V_A(t)\frac{1 + \delta_t V_A(t)}{2} \quad (S2.6)$$

If we substitute (S2.2) and (S2.5) into (S2.6), we get:

$$V_A(t+1) = \frac{V_A(0)}{2} + V_A(t)\left(\frac{1 + \rho_g(t)}{2}\right) \quad (S2.7)$$

This equation matches what is reported in other sources[13-15].

We will also define the intergenerational genetic variance ratio, $Q_A(t)$, as we will refer to that later:

$$Q_A(t) = \frac{V_A(t+1)}{V_A(t)} \quad (S2.8)$$

To see how genetic variance changes across generations under different scenarios, we apply Equation (S2.7) recurrently. In Supplementary Figure 6, we see how the genetic variance ($V_A$) increases in successive generations of assortative mating following Equation (S2.7) for three different phenotypic partner correlations. All three example assume that the heritability is 50% in the base population, and that the genetic variance starts at unit variance $V_A(0) = 1$. Panel A shows how the variance increases asymptotically towards equilibrium (indicated by the dashed grey lines), and Panel B shows the intergeneration variance ratio ($Q_A$). Overall, variance increases in the initial generations of assortment before reaching equilibrium after between 6 to 10 generations, practically speaking, depending on assortment strength. Larger partner correlations lead to more generations in disequilibrium in part because increased genetic variance leads to increased heritability (all else equal), which in turn leads to an increased genotypic correlation between partners, which in turn increases the variance further.

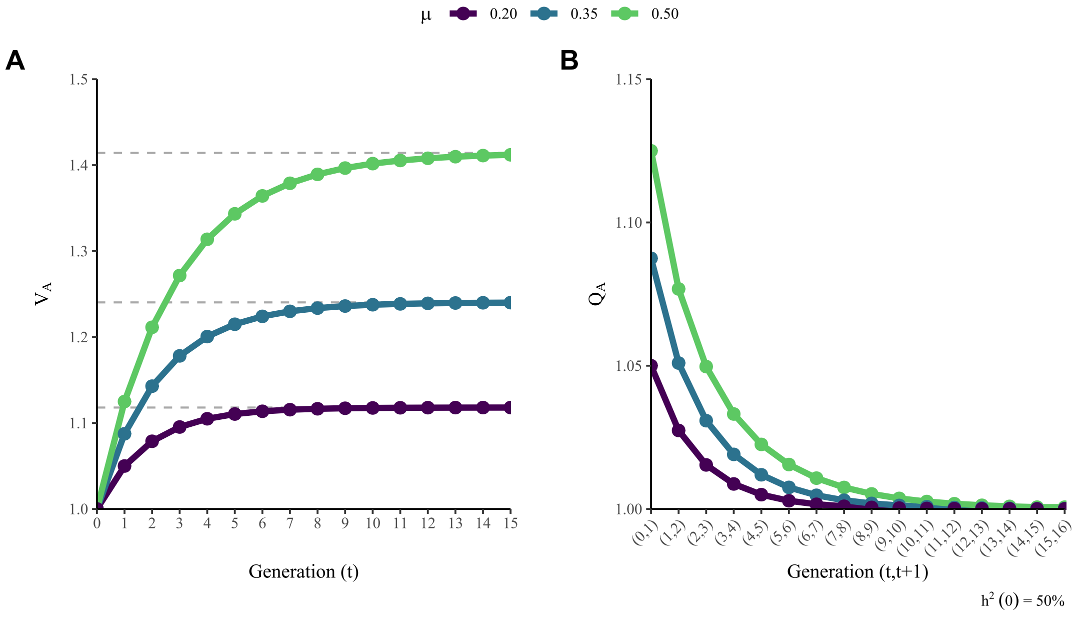

**Supplementary Figure 6** *Genetic variance in successive generations of assortative mating.*

Note that we can simplify Equation (S2.7) further if we are interested in the first generation of assortment, where $V_A(t) = V_A(0)$, or the equilibrium variance, where $V_A(t) = V_A(t+1) = V_A(\infty)$:

$$V_A(1) = V_A(0)\left(1 + \frac{\rho_g(0)}{2}\right) \quad (S2.9a)$$

$$V_A(\infty) = \frac{V_A(0)}{1 - \rho_g(\infty)} \quad (S2.9b)$$

These simplifications match equations reported elsewhere[4,13].

#### 2.2 Gene-environment correlations during disequilibrium

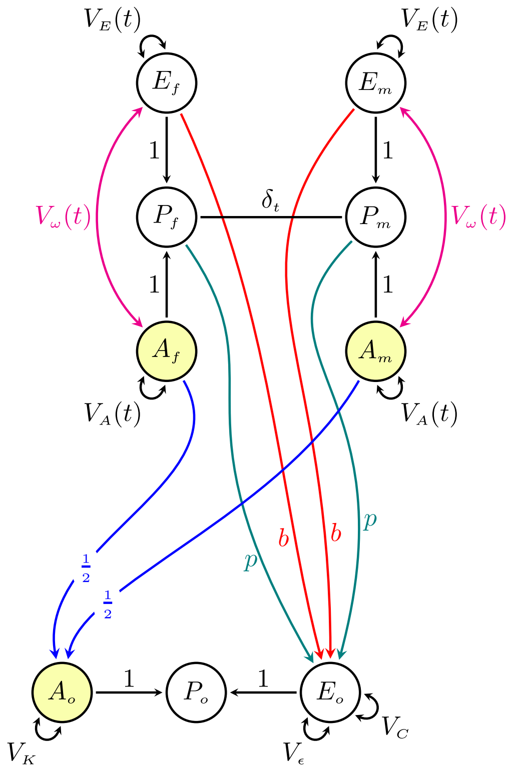

**Supplementary Figure 7** *This path diagram adapts Supplementary Figure 5 by also adding cultural and phenotypic transmission (in addition to genetic transmission). Under disequilibrium, the environmental variance, $V_E(t)$, and gene-environment covariance, $V_\omega(t)$, will change across generations too, necessitating generation-specific notation for these as well.*

In Supplementary Figure 7, the offspring environment are caused by various factors. Most importantly, there are causal effects from the parental environments (denoted $b$, coloured red) and from the parental phenotypes (denoted $p$, coloured teal). Note that these are unstandardized effects and will depend on the variance of the parental environment and phenotype respectively. There are also two sources of environmental variance that are not shared between parents and offspring: $V_C$ indicates environmental variance shared by siblings (not drawn), and $V_\epsilon$ denotes environmental variance unique to the individual. Both of those are for simplicity assumed to be constant across generations.

As we saw in Note 1.2, environmental transmission will induce gene-environment correlations, especially when coupled with assortative mating. Because we have already defined $\omega$ as the gene-environment *correlation*, Supplementary Figure 7 uses $V_\omega(t)$ to denote the gene-environment *covariance* in generation $t$.

##### *Genetic variance*

Supplementary Figure 6 assumed that similarity between offspring was entirely attributed to genetic transmission. What if there were environmental transmission as well? If this is the case, then the equations for the resulting variances and gene-environment correlations become quite complicated, in part because the environmental variance and gene-environment covariance will also change across generations (which is why they have generation-specific notation in Supplementary Figure 7). However, as we shall see, the genetic variance in generation $t + 1$ is still just a function of the genetic variance and the genotypic partner correlation in the preceding generation. In other words, gene-environment correlations will, just as during equilibrium, mimic the effect of higher heritability. Let us start by tracing the genotypic correlation between partners (i.e., the genotypic covariance divided by the genetic variance):

$$\rho_g'(t) = \frac{\delta_t\big(V_A(t) + V_\omega(t)\big)^2}{V_A(t)} \tag{S2.10}$$

We can also trace the genetic variance in the offspring generation. Even though there are many more pathways, when we simplify, we find that it is still just a function of the genetic variance in the preceding generation and the genotypic correlation between partners:

$$V_A(t+1) = Var(A_o)$$

$$= V_K + \frac{V_A(t)}{4} + \frac{V_A(t)}{4} + \frac{\delta_t V_A^2(t)}{4} + \frac{\delta_t V_A^2(t)}{4} + \frac{\delta_t {V_\omega}^2(t)}{4} + \frac{\delta_t {V_\omega}^2(t)}{4} + \frac{V_A(t)\delta_t V_\omega(t)}{4} + \frac{V_\omega(t)\delta_t V_A(t)}{4}$$

$$= V_K + \frac{V_A(t)}{2} + \frac{\delta_t\big(V_A(t) + V_\omega(t)\big)^2}{2}$$

$$= V_K + V_A(t)\frac{1 + \rho_g'(t)}{2} \tag{S2.11}$$

As is apparent, gene-environment correlations do not affect the equations for genetic variance across generations except through its effect on increased genetic similarity between partners. However, it does affect the heritability across generations. To see why, we must find the equations for environmental variance and gene-environment correlations in generation $t + 1$.

##### *Environmental variance*

If environmental transmission results from cultural transmission (denoted $b$), the environmental variance in generation $t + 1$ will depend on the environmental variance in the preceding generation, $V_E(t)$, along with the environmental correlation ($\rho_e'$) between partners. If, on the other hand, it is the result of phenotypic transmission (denoted $p$), then the environmental variance in generation $t + 1$ will depend on the phenotypic variance in the preceding generation, $V_P(t) = Var(P_f) = Var(P_m)$, along with the phenotypic correlation ($\mu$) between partners. When we trace the environmental variance and simplify, we find that:

$$V_E(t+1) = Var(E_o)$$

$$= V_\epsilon + V_C + 2V_E(t)b^2\big(1 + \rho_e'(t)\big) + 2V_P(t)p^2(1+\mu) + 4pb\big(V_E(t) + V_\omega(t)\big)(1+\mu) \tag{S2.12}$$

Where the environmental and phenotypic correlations between partners are:

$$\rho_e'(t) = Corr\big(E_f, E_m\big) = \frac{\delta_t\big(V_E(t) + V_\omega(t)\big)^2}{V_E(t)} \tag{S2.13a}$$

$$\mu = Corr\big(P_f, P_m\big) = \frac{\delta_t\big(V_A(t) + V_E(t) + 2V_\omega(t)\big)^2}{V_A(t) + V_E(t) + 2V_\omega(t)} = \delta_t\big(V_A(t) + V_E(t) + 2V_\omega(t)\big) \tag{S2.13b}$$

##### *Gene-environment correlations*

Assortative mating will also induce and increase the gene-environment correlation. We can trace the gene-environment covariance in Supplementary Figure 7. After simplification, we find that:

$$V_\omega(t+1) = Cov(A_o, E_o)$$

$$= V_\omega(t)b + \big(V_A(t) + V_\omega(t)\big)\delta_t\big(V_E(t) + V_\omega(t)\big)b + \big(V_A(t) + V_\omega(t)\big)(p + p\mu) \tag{S2.14}$$

Because $V_A$ and $V_E$ will increase across generations, so will $V_\omega$. To get the gene-environment *correlation* ($\omega$), we can divide by the geometric mean of the genetic and environmental variance:

$$\omega(t) = Corr(A_f, E_f) = \frac{V_\omega(t)}{\sqrt{V_A(t) \times V_E(t)}} \qquad (S2.15)$$

Even though the denominator increases under assortative mating, the numerator increases more quickly. In other words, the gene-environment correlation will increase in successive generations of assortative mating, all else equal. See the simulation in Note 5.4 for a demonstration.

##### ***Heritability across generations***

Now that we have expressions for all the terms that go into phenotypic variance, we can look at how the heritability changes across generations with assortative mating. The heritability is, as always, the proportion of phenotypic variance attributable to genetic variance.

$$h^2(t+1) = \frac{V_A(t+1)}{V_A(t+1) + V_E(t+1) + 2V_\omega(t+1)} \qquad (S2.16)$$

In the absence of environmental transmission, heritability will increase across generations under assortative mating. However, when there is assortative mating, the other components of the phenotypic variance also increase, which in turn make the changes in heritability less predictable. Because $V_E + 2V_\omega$ can increase faster than $V_A$, the heritability may decrease or remain constant across generations, even though the genetic variance increases. See the simulation in Note 5.4 for a demonstration.

#### 2.3 The genotypic correlation between parents and offspring under disequilibrium

To find the genotypic correlation between parents and offspring, we must know the variance (which we just derived) and the covariance. To find the covariance, we apply path tracing rules to find all valid chains between $A_m$ (or $A_f$) and $A_o$ in Supplementary Figure 7, and simplify accordingly:

$$\begin{aligned} Cov(A_m, A_o) &= \frac{V_A(t)}{2} + \frac{\delta_t\big(V_A(t) + V_\omega(t)\big)^2}{2} \\ &= V_A(t)\left(\frac{1 + \rho_g'(t)}{2}\right) \end{aligned} \quad (S2.17)$$

Because the genotypic covariance is a function of the genetic variance in the parent generation, the genotypic covariance too will increase together with the variance in successive generations of assortative mating. To find the correlation, we divide the covariance by the geometric mean of the variances:

$$r_{g_{PO}}'(t) = Corr(A_m, A_o) = \frac{V_A(t)\left(\frac{1 + \rho_g'(t)}{2}\right)}{\sqrt{V_A(t)V_A(t+1)}} \quad (S2.18)$$

The purple line Supplementary Figure 8A shows how the genotypic correlation between parents and offspring increases with successive generations of assortative mating (assuming $h^2(0) = 50\%$, $\omega = 0$, and $\mu = .50$). Note that, because the parent-offspring correlation is an intergenerational correlation, the number of prior generations of assortative mating will not be the same for parents and offspring. We will keep denoting the parental generation as generation $t$ (and therefore the offspring generation as $t + 1$). We see that the correlation increases immediately and keeps increasing towards an equilibrium just like how the variance increases.

The green dotted line is the expected genotypic correlation if we assumed equilibrium and used Equation (2) from the main paper ($r_{g_1} = \frac{1+\rho_g}{2}$). (Remember, Equation (2) and Equation (5) are equal for first-degree relatives). We see that it only matches the actual correlation when the trait is in equilibrium. Otherwise, it overestimates the correlation. A mismatch between the true correlation between relatives and expected correlation given Equation (5) is therefore evidence of a trait in disequilibrium.

The reason for this mismatch is directly attributable to different variances across generations. Equation two in the main paper implicitly assumes that the variance is equal across generations, which cancels out the variance terms in the denominator and numerator. However, during disequilibrium, the geometric mean of the variances (the denominator) is slightly more than the variance in the parent generation (part of the numerator). This leads to a smaller correlation than Equation (2) would predict. When we rearrange Equation (S2.8) and substitute that into the denominator of Equation (S2.18), the variance terms cancel out and we can express the correlation as:

$$r_{g_{PO}}(t) = \frac{1 + \rho_g'(t)}{2\sqrt{Q(t)}} \quad (S2.19)$$

The *degree* of mismatch depends on the number of generations since assortative mating began. It should therefore be possible to infer something about the history of assortment by quantifying the degree of mismatch in the parent-offspring correlation. Torvik, et al. [17] defined $U$ as the ratio of observed increase in correlation to the expected increase in correlation at equilibrium given the current genotypic correlation between partners. In other words, it is the observed increase as a percentage of the expected increase at equilibrium, where 100% would indicate equilibrium:

$$U(t) = \frac{Actual\ Increase}{Expected\ Increase} = \frac{r_{g_{PO}}(t) - 0.5}{\left(\frac{1+\rho_g'(t)}{2}\right) - 0.5} \quad (S2.20)$$

Note that $U$ is specific to each relationship and will not be the same for full siblings or other relatives. Here, we are using it for the genotypic parent-offspring correlation. Panel B in Supplementary Figure 8 plots how $U$ develops in successive generations of assortative mating depending on the phenotypic correlation between partners, starting from a random mating population. We see that it is approximately 72% after one generation, 84% after two generations, 90% after three generations, etc., For traits under stronger assortment, $U$ increases slightly slower (and vice versa), but the differences are not large.

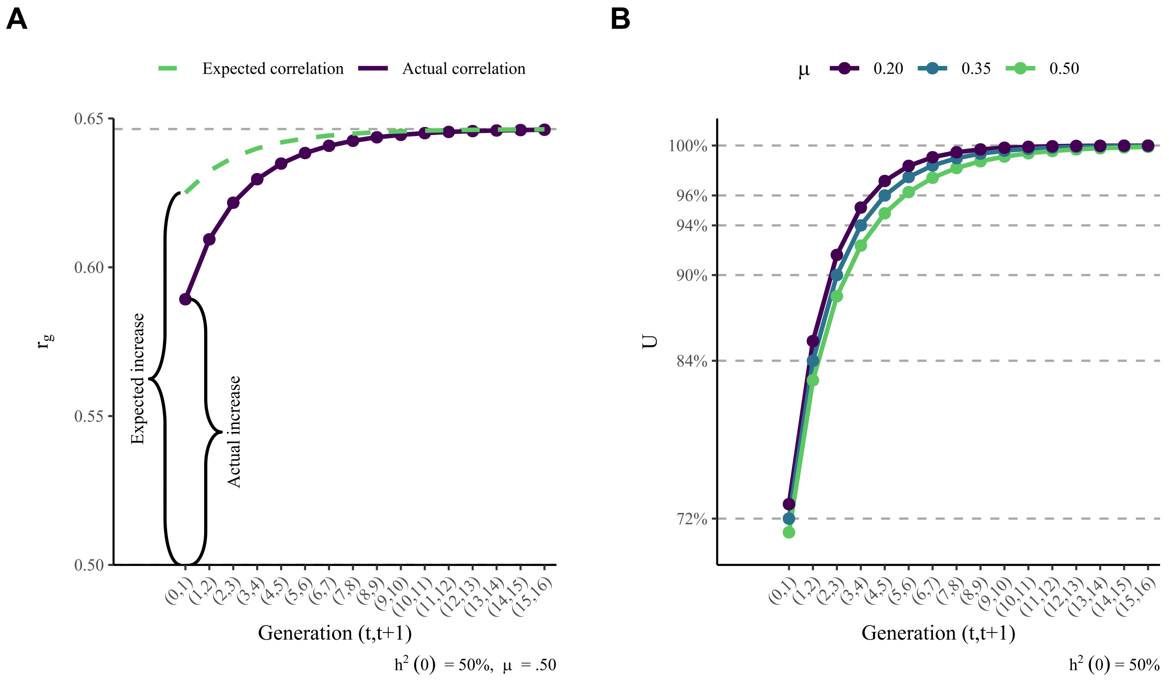

**Supplementary Figure 8** *The dynamics of the genotypic parent-offspring correlation during disequilibrium.*

By comparing an observed $U$-value with Supplementary Figure 8B, it is possible to infer the equivalent numbers of generations since assortative mating began. Obviously, this would involve making assumptions such as constant environmental variance and constant partner correlations (preceded by random mating), which is unlikely to reflect reality. For this reason, we do not deem it worthwhile to develop a more accurate estimation of $t$ beyond looking up a rough estimate in the figure (although it is perfectly possible to calculate $U(t)$ with other assumptions). A given $U$ can be said to be *equivalent* to an approximate $t$, which is to say that a population with no assortative mating prior to $t$ generations of constant, univariate assortment would yield a similar $U$ value. It is important to note, though, that multiple other processes could give rise to the same $U$ value, such as changes in assortment strength, changes in environmental and genetic effects across generations, changes in gene-environment correlations, or possibly changing patterns of assortative mating across traits.

Note that $U$ calculated with polygenic indices roughly tracks the true genotypic partner correlation of the sorted phenotype, not the polygenic index correlation between partners (which is confounded by the genetic signal, see Note 3.4). Given that educational attainment appears to be roughly 50% heritable[18,19] and exhibits a strong phenotypic correlation between partners[17,20], we can search for its observed U-value ($U = 90\%$) along the green line in Supplementary Figure 8B. We find the U-value roughly matches what we would expect at $t = 2$, indicating two prior generations of assortment. In other words, our result for educational attainment matches what we would expect if the parental generation was the *third* generation to mate assortatively on educational attainment.

##### *A brief note on siblings*

If you were to expand Supplementary Figure 7 to include siblings in the offspring generation and trace the genotypic covariance, you would find that it equals the genotypic covariance between parent and offspring. The correlation, on the other hand, would be slightly different because the denominator would now be the variance in the offspring generation only, $V_A(t+1)$, instead of the geometric mean of the variances in the two generations, $\sqrt{V_A(t)V_A(t+1)}$. Because the offspring variance is bigger than the parent variance during disequilibrium, the denominator would be larger, and the resulting correlation would be smaller. This, in turn, means that the associated $U$-values in any given $t$ will be slightly less for sibling correlations compared to parent-offspring correlations. Nevertheless, as you can see in the simulations in Supplementary Note 5, there are no substantive differences in when they reach equilibrium.

### Supplementary Note 3 Polygenic indices

In this note, we describe polygenic index correlations between family members under assortative mating. In Note 3.1, we describe polygenic index correlations at equilibrium in a model with genetic transmission only, showing that the polygenic index correlation will be attenuated depending on the correlation between the polygenic index and the true genetic factor. In Note 3.2, we expand this model with environmental transmission, showing that this not affect the polygenic index correlations beyond its effect on the true genotypic correlation. In the final two subsections, we describe polygenic indices at disequilibrium. In Note 3.3, we describe the polygenic index variance across generations, and in Note 3.4, we describe the polygenic index correlation between parents and offspring during disequilibrium.

##### *Key takeaways*

- Assortative mating's effect on polygenic index correlations will depend on the squared correlation between the polygenic index and the true genetic factor (i.e., the genetic signal).
- The polygenic index correlation between any two family members will be biased towards the coefficient of relatedness if the genetic signal is low.
- At equilibrium, the relationship between the polygenic index correlation between partners and the polygenic index correlation between *first-degree* relatives is the same regardless of genetic signal. For higher degree relatives, the true polygenic index correlation will, depending on the genetic signal, be *slightly* higher than Equation (5) would predict based on the polygenic index correlation between partners.
- Gene-environment correlations will affect the correlation between the genotype and phenotype, which affects the genetic consequences of assortative mating (as described in Note 1.2). However, gene-environment correlations will not have any additional effects on polygenic index correlations beyond their effect on true genotypic correlations (where it mimics higher heritability).
- Because genetic variance increases during disequilibrium, so does the polygenic index variance (albeit to a lesser extent, depending on the genetic signal).
- The increase in polygenic index variance increases the genetic signal until it reaches an equilibrium (similar to how heritability increases with assortative mating). This means that the genetic signal is not merely the correlation between true direct effects and the respective weight for the polygenic index.
- The polygenic index correlation between parents and offspring increases with successive generations of assortative mating (just like the true genotypic correlation).
- The degree of mismatch between the correlation predicted by Equation (5) and the true polygenic index correlation is almost unaffected by the strength of the genetic signal. This means that the degree of mismatch is informative on deviations from equilibrium and thereby the history of assortative mating without making strong assumptions about the genetic signal (although one must make assumptions on the true heritability and assortment strength).

#### 3.1 Polygenic index correlations

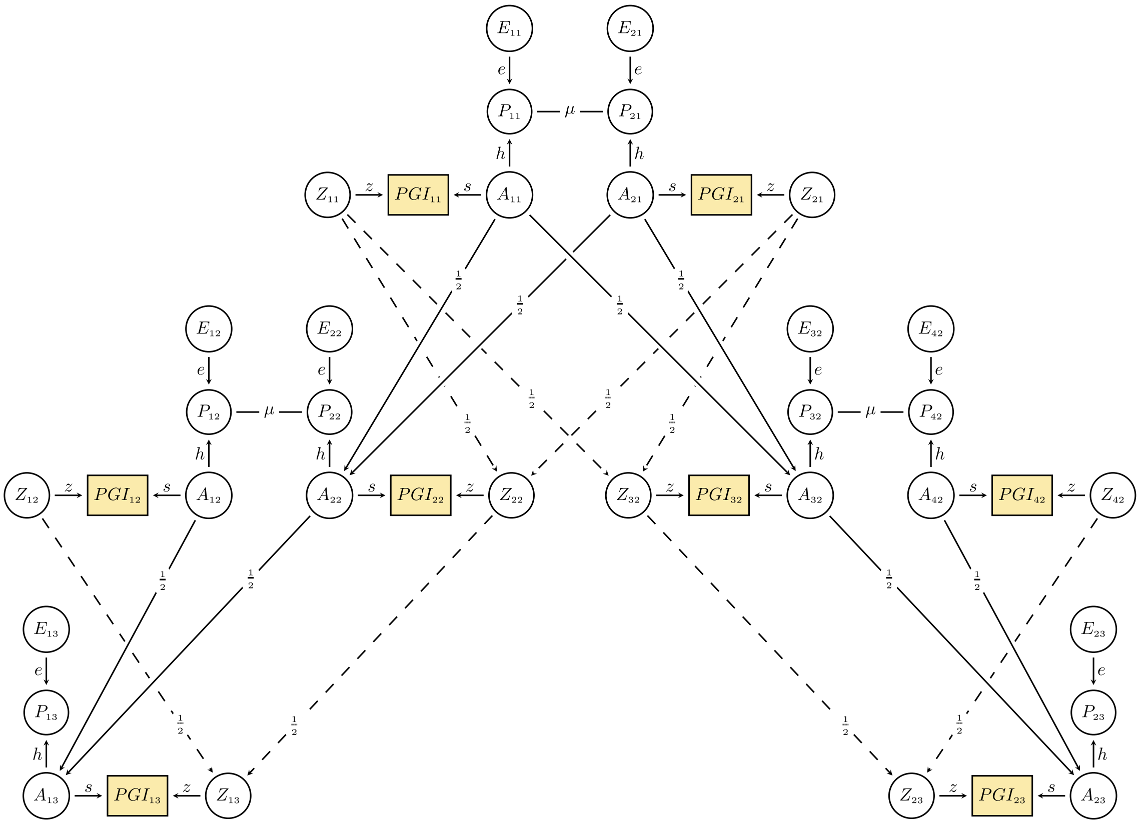

**Supplementary Figure 9** *Path diagram of associations between relatives' polygenic indices (PGI).*

Supplementary Figure 9 is an extension of Fig. 1 and represents the associations between various family members' polygenic indices at equilibrium. The equilibrium assumption is instantiated via equal variances across generations. All variables have unit variances, and the variance terms (↔) are excluded to remove clutter. We assume polygenic indices are composed of true genetic factors ($A$) and noise ($Z$). That is, each loci have an effect on the polygenic index (i.e., the *weight*), which, depending on its accuracy, differ from its effect on the phenotype. Note that many different causes of polygenic index inaccuracy (such as including loci without true effects, missing loci that have true effects, or overestimating the loci's true effects due to linkage disequilibrium) all amounts to the same thing: A difference between the loci's true effects on the phenotype and on the polygenic index. It is therefore not necessary to distinguish between these sources of polygenic index inaccuracy in the path diagram and in the following derivations.

If you sum the effects of all loci (including their covariance), you are left with the total covariance between the true genetic factors and polygenic index. This can, when standardized, be summarized as a correlation denoted $s$ in Supplementary Figure 9. This, in turn, means that the genetic signal – the shared variance between the polygenic index and the true genetic factor – is $s^2$. Just like with $h^2$ (the heritability), $s^2$ will include the covariance between different loci (linkage disequilibrium) and is therefore not just the sum of the individual loci's effects. This is especially true for polygenic indices where the weights themselves are biased by linkage disequilibrium[21]. Instead, the genetic signal is the proportion of variation in the polygenic index that can be attributed to the true genetic factor (see also Note 4.4).

To keep all variables at unit variances, the effect of noise will be $z = \sqrt{1 - s^2}$ (similar to how $e = \sqrt{1 - h^2}$) and have a higher value when $s$ is low. Because the error in the polygenic index primarily results from misestimated effects and not mismeasured genotypes, the errors will be inherited just like the true genetic factors. Using path tracing, we see that the polygenic index correlation between partners (e.g., $PGI_{11} \leftrightarrow PGI_{21}$) should be:

$$\rho_{pgi} = s \times h \times \mu \times h \times s = \mu h^2 s^2 \qquad (S3.1)$$

If one suspects that the genetic signal is poor, it is possible to do a back-of-the-envelope calculation to find the approximate value of $\rho_g$ (the true genotypic partner correlation). For example, the polygenic index from the most recent genome-wide association study for educational attainment explained between 12% and 15% of the variance in holdout samples[22], whereas a meta-analysis of twin studies on educational attainment[19] estimated the mean heritability to be about 45%. Obviously, these values are likely to vary by population and cohort, and may depend on violated assumptions, but if we take these at face value, the polygenic index captures $s^2 = \frac{15}{45} = 33\%$ of the true additive genetic factor. We observe a polygenic index correlation of .14 between partners for educational attainment, but if that polygenic index has a genetic signal of 33%, then the true genotypic correlation would be somewhere around $\rho_g = \frac{\rho_{pgi}}{s^2} =.42$. (Remember that the confidence intervals would also be similarly inflated). If you think the genetic signal is higher or lower, you can repeat the calculation with numbers of your choice.

The polygenic index correlation between any two relatives can be thought to be composed of two components: the error-related genetic factor and the phenotype-related genetic factor. The error term correlation should equal the coefficient of relatedness, whereas the phenotype-related genotypic correlation will equal the true genotypic correlation. These two components will be weighted by $z^2$ and $s^2$, respectively:

$$r_{pgi_k} = z^2 \left(\frac{1}{2}\right)^k + s^2 \left(\frac{1+\rho_g}{2}\right)^k \qquad (S3.2)$$

This can be verified by tracing all chains between any two relatives in Supplementary Figure 9. For first degree-relatives ($k = 1$), this simplifies to $\frac{1+\rho_{pgi}}{2}$. For higher degree relatives, the exact correlation will depend on the genetic signal. It can be approximated with $r_{pgi_k} \approx \left(\frac{1+\rho_{pgi}}{2}\right)^k$, although this will *slightly* underestimate the correlation (see Supplementary Figure 10). The reason for this is that the genetic signal must only be included once regardless of relation, whereas $r_{pgi_k} \approx \left(\frac{1+\rho_{pgi}}{2}\right)^k$ will include it once *for each* degree of relatedness. That is, it will raise the genetic signal to the degree of relatedness ($s^{2k}$), thus biasing the approximation towards the degree of relatedness.

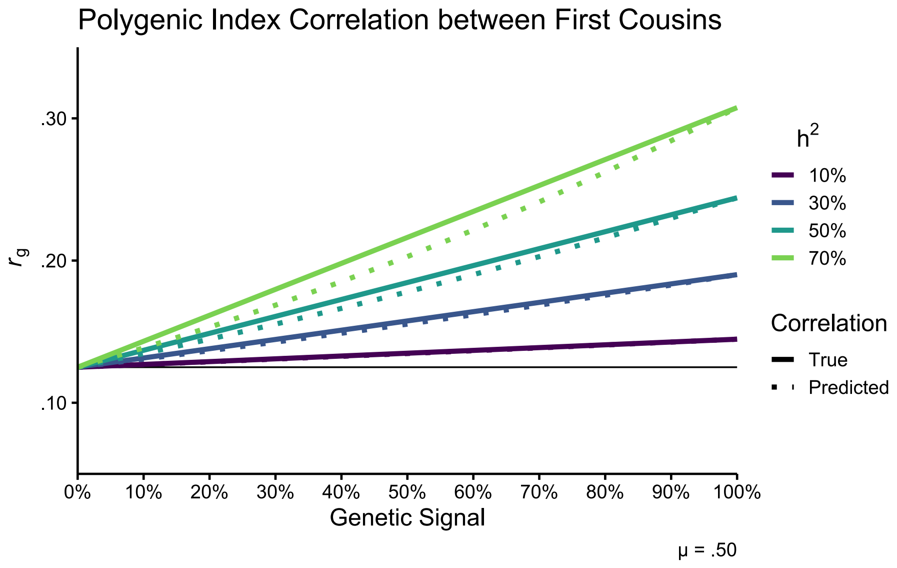

**Supplementary Figure 10** *Polygenic index correlations between first cousins under various genetic signals. The dotted lines are the predicted correlation using the polygenic index correlation between partners (i.e., Equation (5)).*

#### 3.2 Polygenic index correlations with gene-environment correlations

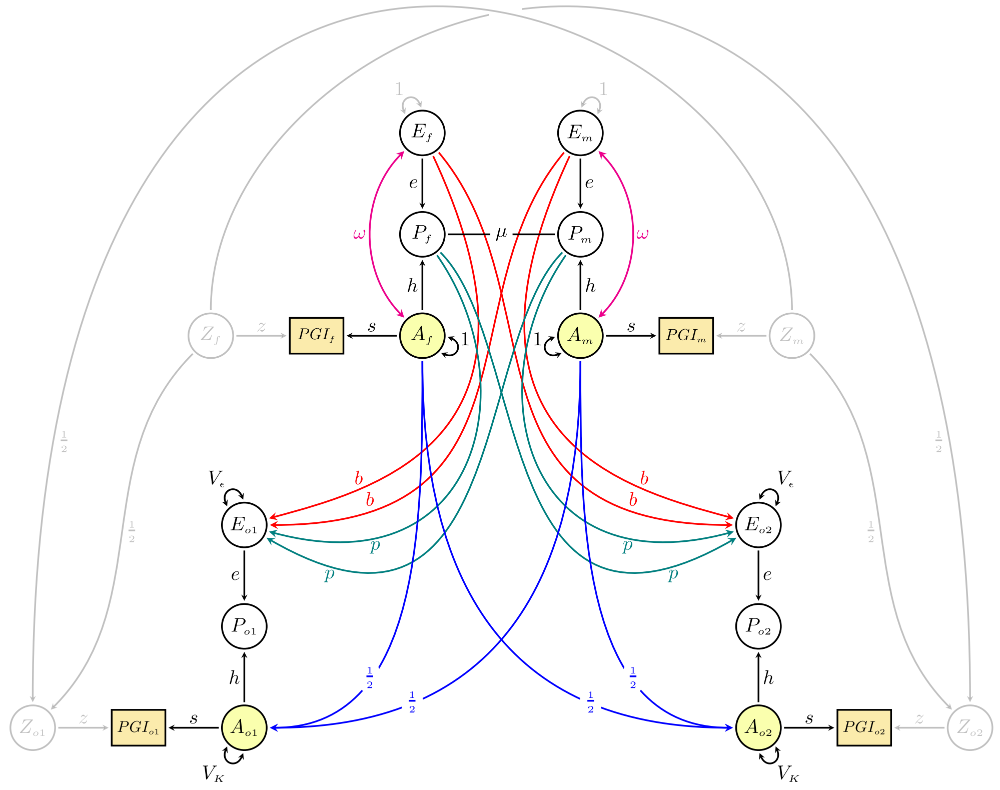

**Supplementary Figure 11** *Associations between relatives' polygenic indices (PGI) under gene-environment correlations.*

The key difference between the path diagram in Supplementary Figure 11 compared to Supplementary Figure 9 is that the correlation between the polygenic index and the phenotype now includes a gene-environment correlation. The correlation between, for example, $PGI_f$ and $P_f$ is no longer only the direct effect of genetic differences ($sh$), but also the indirect association through the gene-environment correlation ($s\omega e$). Just like with the true genetic factor, this extra pathway will increase the correlation between the polygenic indices of partners. If we trace $PGI_f \leftrightarrow PGI_m$, we find:

$$\rho'_{pgi} = \mu(h + \omega e)^2 s^2 \tag{S3.3}$$

When we trace the correlation between the polygenic indices of siblings ($PGI_{o1} \leftrightarrow PGI_{o2}$) or parent-offspring (e.g., $PGI_f \leftrightarrow PGI_{o1}$), we find that there are no new pathways other than those between the genotypes of partners:

$$r'_{pgi_1} = z^2\left(\frac{1}{2}\right) + s^2\left(\frac{1+\rho'_g}{2}\right) = \frac{1+\rho'_{pgi}}{2} \tag{S3.4}$$

Hence, the polygenic index correlation for first-degree relatives is equally proportional to the polygenic index correlation between partners, regardless of gene-environment correlations or genetic signal. We have already established that this is not the case for higher-degree relatives, whose correlation depend on the both the degree of gene-environment correlation and the genetic signal. If you were to expand Supplementary Figure 11 to include higher-degree relatives, you would find that the polygenic index correlation would be the true genotypic correlation (as derived in Note 1.2) multiplied by the genetic signal. They can still be approximated by $r'_{pgi_k} = \left(\frac{1+\rho'_{pgi}}{2}\right)^k$, although this will underestimate the true correlation somewhat if there are substantial gene-environment correlations (as explained in Note 1.2) or if the genetic signal is poor (as explained in Note 3.1).

##### 3.3 Polygenic index variance during disequilibrium

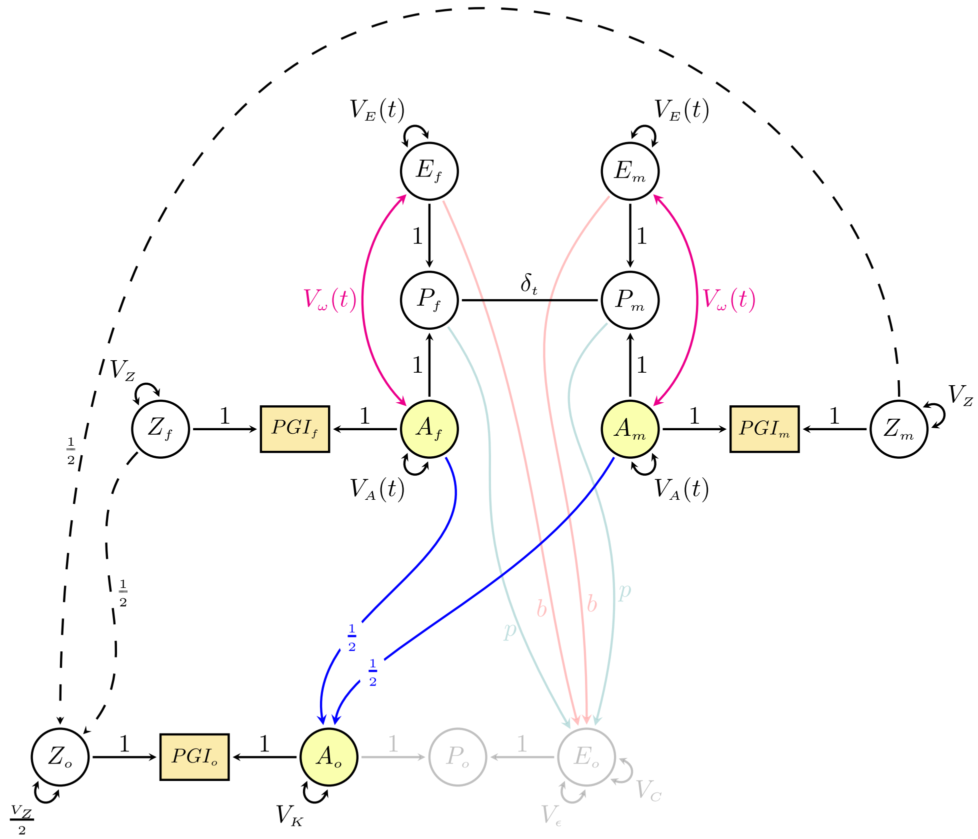

**Supplementary Figure 12** *Path diagram illustrating polygenic index (co)variance under disequilibrium. Pathways that are irrelevant for understanding polygenic index similarity among the depicted individuals are greyed out to remove clutter.*

Supplementary Figure 12 extends Supplementary Figure 6 by including polygenic indices. Because we are no longer assuming unit variances, the genetic signal is now indicated by the relative importance of true genetic variance, $V_A(t)$, and error variance, $V_Z$ in the polygenic index. Because the initial variance in the polygenic index is arbitrary, we will for simplicity's sake not scale the variance (without loss of generality). Applying path tracing rules, we find that the variance in the polygenic index is:

$$V_{pgi}(t) = Var(PGS_m) = V_A(t) + V_Z \tag{S3.5}$$

Because genetic variance initially increases with successive generations of assortative mating, the variance in the polygenic index also increases. This has the slightly odd effect of making the genetic signal higher in successive generations. We must therefore give the genetic signal generation-specific notation:

$$s^2(t) = \frac{V_A(t)}{V_A(t) + V_Z} \tag{S3.6}$$

When using polygenic indices as proxies for the true genetic factor, we must understand how the genetic signal affects $Q$ and $U$. We will consider $U$ in the next section. Let us first consider how the polygenic index variance changes across generations. Because we already know how the genetic variance changes, we can combine Equation (S3.5) and (S2.7):

$$V_{pgi}(t+1) = V_A(t+1) + V_Z \tag{S3.7}$$

Likewise, we change adapt Equation (S2.8) and obtain the intergenerational variance ratio for the polygenic index:

$$Q_{pgi}(t) = \frac{V_{pgi}(t+1)}{V_{pgi}(t)} \tag{S3.8}$$

Because the polygenic index variance contains noise that presumably stays constant across generations, the intergenerational variance ratio for polygenic indices will always be closer to one than the true variance ratio, meaning it will underestimate the increased variance in the offspring generation under positive assortative mating:

$$|1 - Q_{pgi}| \leq |1 - Q_A| \tag{S3.9}$$

Put another way, the polygenic index variance will not increase to the same extent as the true genetic variance. If the genetic signal is known (or we are willing to assume a given signal), then it is possible to calculate the true intergenerational variance ratio ($Q_A$) given the variance ratio for polygenic indices ($Q_{pgi}$). Before that, we must do some rearranging. To declutter the following equations, we will not use the generation-specific notation if the parameter is from generation $t$.

First, we rearrange Equation (S3.6) to give us the polygenic variance attributable to noise ($V_Z$) using $s^2$:

$$V_Z = V_A \frac{(1 - s^2)}{s^2} \tag{S3.10}$$

We then rearrange Equation (S2.8) to give us $V_A(t+1)$ using $Q_A$:

$$V_A(t+1) = Q_A V_A \tag{S3.11}$$

We can now substitute (S3.10) and (S3.11) into Equations (S3.5) and (S3.7) and simplify accordingly:

$$\begin{aligned} V_{pgi} &= V_A + V_A \frac{(1 - s^2)}{s^2} \\ &= \frac{V_A}{s^2} \end{aligned} \tag{S3.12a}$$

$$\begin{aligned} V_{pgi}(t+1) &= Q_A V_A + V_A \frac{(1 - s^2)}{s^2} \\ &= V_A\left(Q_A + \frac{1}{s^2} - 1\right) \end{aligned} \tag{S3.12b}$$

Which we in turn can substitute into (S3.8) and simplify accordingly:

$$\begin{aligned} Q_{pgi} &= \frac{V_A(Q_A + 1/s^2 - 1)}{V_A/s^2} \\ &= Q_A s^2 + 1 - s^2 \end{aligned} \tag{S3.13}$$

Solving for $Q_A$ gives us:

$$Q_A(t) = 1 + \frac{Q_{pgi}(t) - 1}{s^2(t)} \tag{S3.14}$$

We can continue using educational attainment as an example, where the polygenic index variance was $Q_{pgi} = 1.023$ times greater in the offspring generation than in the parental generation. If we keep assuming the genetic signal is 33.3% (see Note 3.1), the true genetic factor has 1.069 times (or 6.9%) greater variance in the offspring generation than the parent generation.

#### 3.4 The polygenic index correlation between parents and offspring under disequilibrium

The polygenic index covariance between parents and offspring consists of two sets of pathways: The true genotypic covariance (derived in Note 2.3) and the covariance attributable to the shared error terms. The polygenic index covariance is simply the sum of these two components:

$$Cov(PGI_m, PGI_o) = V_A(t)\left(\frac{1+\rho_g'(t)}{2}\right) + \frac{V_Z}{2} \qquad (S3.15)$$

You can double-check this by tracing all valid chains in Supplementary Figure 12. To get the correlation, we must divide by the pooled variance:

$$r_{pgi}(t) = Corr(PGI_m, PGI_o) = \frac{V_A(t)\left(\frac{1+\rho_g'(t)}{2}\right) + \frac{V_Z}{2}}{\sqrt{V_{pgi}(t)V_{pgi}(t+1)}} \qquad (S3.16)$$

Because the genetic signal changes from generation to generation, it is not straightforward to reduce (S3.15) or (S3.16) to simpler equations (nor is it necessary here). Instead, we can calculate what the observed correlation would be under various assumptions and see how it behaves compared to the true genotypic correlation. It will also be useful to see how it behaves compared to the expected correlation given the polygenic index correlation between partners (i.e., $\frac{1+\rho_{pgi}}{2}$). The polygenic index correlation between partners will still equal Equation (S3.1) (i.e., $\rho_{pgi}' = \rho_g' s^2$), which you can double check by tracing all valid chains between $PGS_f$ and $PGS_m$ in Supplementary Figure 12 and divide by the polygenic index variance:

$$\rho_{pgi}'(t) = \frac{\delta_t\big(V_A(t) + V_\omega(t)\big)^2}{V_A(t)/s^2(t)} = \frac{s^2(t)\delta_t\big(V_A(t) + V_\omega(t)\big)^2}{V_A(t)} = \rho_g'(t)s^2(t) \qquad (S3.17)$$

We can also define the polygenic index equivalent of $U$ (i.e., the ratio of actual to expected increase):

$$U_{pgi}(t) = \frac{r_{pgi}'(t) - 0.5}{\left(\frac{1+\rho_{pgi}'(t)}{2}\right) - 0.5} \qquad (S3.18)$$

Supplementary Figure 13A shows how various degrees of genetic signal ($s^2$) affects the polygenic index correlation between parents and offspring in successive generations of assortative mating (assuming $h^2(0) = 50\%, \omega = 0$, and $\mu = .50$). The solid lines are the true correlations, whereas the dotted lines are what you would predict using $\frac{1+\rho_{pgi}'(t)}{2}$. We see that less genetic signal (i.e., lower $s^2$) results in a lower polygenic index correlation between parents and offspring. However, we see in Panel B that $U_{pgi}$ is almost unaffected by the genetic signal. There is a small change where lower $s^2$ results in slightly higher $U_{pgi}$, but overall, $U_{pgi}$ will approximately equal $U$. An important consideration, therefore, is that $U_{pgi}$ depends on the true genotypic partner correlation, not the polygenic index correlation. The latter will be confounded by $s^2$, meaning one cannot use the observed correlation to infer how many generations of assortment has occurred. In Supplementary Figure 14, we show two scenarios that would be indistinguishable if one only had access to polygenic index correlations ($\rho_{pgi} = .157, r_{pgi} = .575, U_{pgi} = 95\%$). Without making assumptions about the true heritability and partner correlation, it would be impossible to say whether this indicated three or four generations since assortment began. To infer $t$ based on $U$, one must therefore make assumptions about the likely true heritability and true assortment strength and look at the relevant line in Supplementary Figure 8B (or calculate it manually).

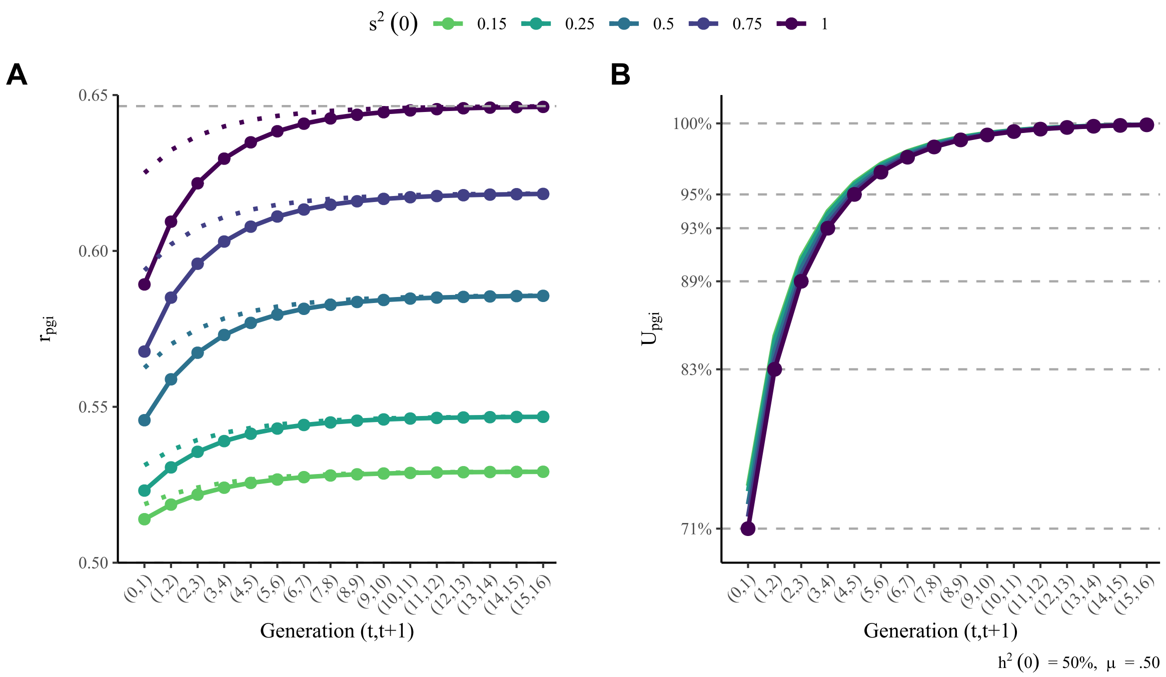

**Supplementary Figure 13** *Dynamics of polygenic score correlations with varying genetic signal strengths*

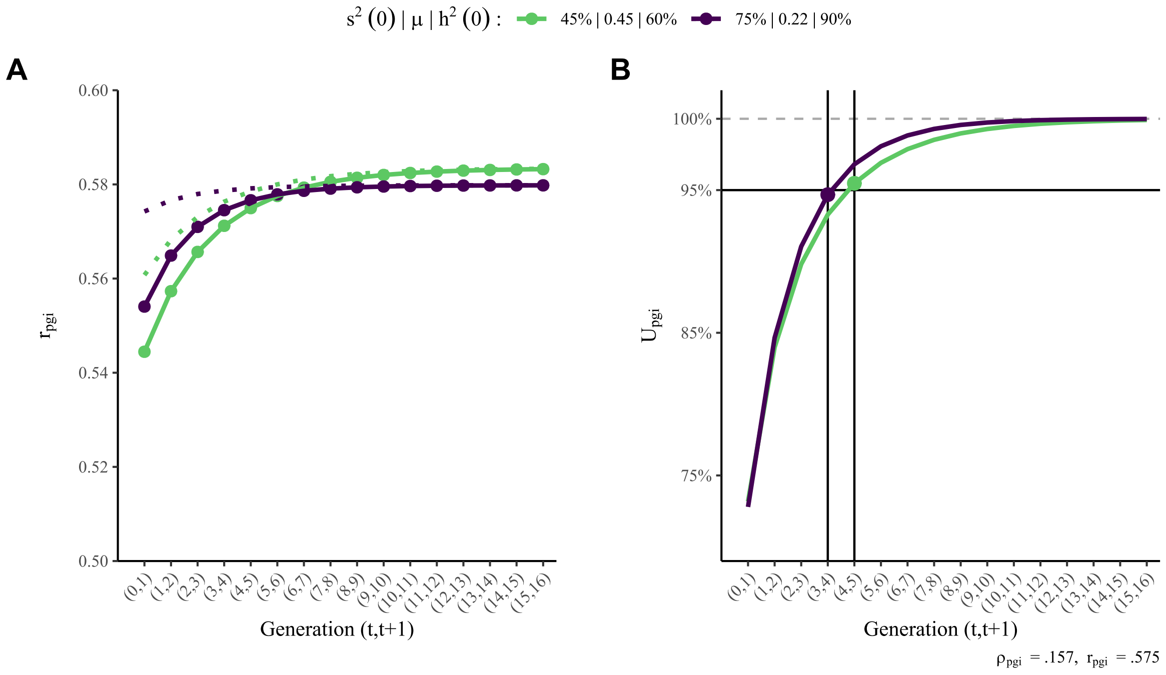

**Supplementary Figure 14** *Similar polygenic index correlations can result from different processes.*

### Supplementary Note 4 Simulations: Method

We started by simulating $2 \times n_{pairs}$ individuals in a randomly mating founder population (generation $t = -2$), which would be the generation before the base population (generation $t = -1$). We did this to accurately simulate siblings in the base population, as well as to be able to calculate a parent-offspring correlation and a first cousin correlation before assortment began. For each new generation after this (i.e., from generation $t = 0$ and onwards), we sorted the would-be partners into pairs based on their phenotypic value, with each pair producing two new offspring. This ensured that that the population size remained constant. The simulation script was written in R and is available at https://osf.io/dgw4r/.

#### 4.1 The genotypes

In the founder population, we randomly generated individual genotypes by simulating $n_{loci}$ loci spread over $n_{chr}$ pairs of chromosomes for each individual $i$. Each locus $l$ consisted of a pair ($f_l$ and $m_l$) of Bernoulli-distributed alleles with a set minor allele frequency ($q_l$), centred to have a mean of 0:

$$f_{li} \sim \text{Bernoulli}(q_l) - q_l$$

$$m_{li} \sim \text{Bernoulli}(q_l) - q_l \qquad (S4.1)$$

Genetic transmission was simulated by mimicking meiosis: A random crossover point was picked for each pair of chromosomes. A new chromosome was then formed by taking the alleles from the start of the first chromosome up to the crossover point and combining them with the alleles from the second chromosome from that point and onwards. This was then paired with the corresponding new chromosome from the other parent. This was repeated for each chromosome, and again for each new individual (with crossover points decided independently and randomly for everyone). Meiosis will automatically create recombination variance, approximately equal to half of the genetic variance in the base population.

#### 4.2 The phenotypes

Each individual $i$ had a phenotype ($P_i$), which was the sum of additive ($A_i$) and dominant ($D_i$) genetic influences and environmental influences ($E_i$):

$$P_i = A_i + D_i + E_i \qquad (S4.2)$$

Each loci's additive effect on the phenotype ($a_l$) was normally distributed with a mean of 0 and a variance equal to $\frac{V_A(0)}{\sum_{l=1}^{n_{loci}} 2q_l(1-q_l)}$, where $V_A(0)$ was the intended additive genetic variance in the base population. The total additive genetic factor was therefore:

$$A_i = \sum_{l=1}^{n_{loci}} a_l \left(f_{l_i} + m_{l_i}\right) \qquad (S4.3)$$

Similarly, the effects of heterozygosity (i.e., dominance deviation) at each loci, $d_l$, was normally distributed with a mean of 0 and a variance equal to $\frac{V_D}{\sum_{l=1}^{n_{loci}} q_l(1-q_l)}$, where $V_D$ is the intended dominant genetic variance in the base population. The total dominant genetic factor was therefore:

$$D_i = \sum_{l=1}^{n_{loci}} d_l \, \mathrm{I}\left(f_{l_i} \neq m_{l_i}\right) \qquad (S4.4)$$

where $\mathrm{I}(E)$ is an indicator function equivalent to wrapping **`as.numeric()`** around a Boolean value in R:

$$\mathrm{I}(E)\begin{cases}1\ if\ E\ is\ true\\0\ if\ E\ is\ false\end{cases}$$

Environmental influences were simulated by letting $E_{ij}$ of individual $i$ in nuclear family $j$ be a function of the phenotypes and environmental factors of the father ($f_j$) and mother ($m_j$), in addition to generation specific environments shared by siblings ($C_j$) and unique to the individual ($\epsilon_{ij}$):

$$E_{ij} = pP_{f_j} + pP_{m_j} + bE_{f_j} + bE_{m_j} + C_j + \epsilon_{ij} \qquad (S4.5a)$$

$$C_j \sim \mathcal{N}(0, V_C) \qquad (S4.5b)$$

$$\epsilon_{ij} \sim \mathcal{N}(0, V_\epsilon) \qquad (S4.5c)$$

Environmental influences in the founder population were simulated as a normally distributed variable ($E$) with a mean of 0 and a variance of $V_E(0)$, where $V_E(0)$ was the intended environmental variance in the base population. If there was need to simulate a pre-existing gene-environment correlation in the founder population, then E was simulated to partially be a function of the genetic influences:

$$E_i = \sqrt{V_E(0)}\left(\omega(0)\frac{A_i}{\sqrt{V_A(0)}} + \sim\mathcal{N}(0{,}1-\omega(0)^2)\right) \qquad (S4.6)$$

#### 4.3 Sorting partners

To attain a given partner correlation, $\mu$, we simulated a sorting variable ($S_i$) which was a function of the standardized phenotype and random error:

$$S_i = \mu\frac{P_i}{\sqrt{Var(P)}} + \mathcal{N}(0{,}1-\mu^2) \qquad (S4.7)$$

Before sorting, individuals were randomly allocated to be either mothers or fathers for the next generation. To prevent accidently simulating inbreeding, full siblings were always the same sex (i.e., allocated to have the same role in the next generation) and therefore not eligible to mate. Would-be mothers and fathers were then separately rank-ordered by $S$, and individuals with the same resulting rank order were considered pairs and were used to produce two offspring for the next generation.

#### 4.4 Simulating the polygenic index

For each individual $i$, we also calculated a polygenic index ($PGI_i$). Each loci's effect on the polygenic index, $s_l$, was the sum of the true genetic effect, $a_l$, and noise, $z_l$, rescaled so that the polygenic index had unit variance in the base population:

$$PGI_i = \sum_{l=1}^{n_{loci}} s_l \left(f_{l_i} + m_{l_i}\right) \quad (S4.8a)$$

$$s_l = \frac{a_l + z_l}{\sqrt{V_A(0) + V_Z}} \quad (S4.8b)$$

$$z_l \sim \mathcal{N}\left(0, \frac{V_Z}{\sum_{l=1}^{n_{loci}} 2q_l(1-q_l)}\right) \quad (S4.8c)$$

$$V_Z = V_A(\infty)\frac{1 - s^2(\infty)}{s^2(\infty)} \quad (S4.8d)$$

where $s^2(\infty)$ was the desired genetic signal in the equilibrium population (i.e., the shared variance between the polygenic index and the true genetic factor). To achieve the desired genetic signal, we adjusted the last values of $z$ (the noise) so that the correlation between the true genetic effects and the polygenic index weights was excatly $s(0)$ (i.e., so it was not subject to sampling variation). In the base population, the correlation between the true direct effects and polygenic index weghts will equal the correlation between the true genetic factor and the polygenic index (i.e., the genetic signal, see Supplementary Figure 15). Under assortative mating, trait-increasing alleles will co-occour in the same individuals, resulting in a larger correlation between the polygenic index and true genetic factor (i.e., a larger genetic signal, see Note 3.3).

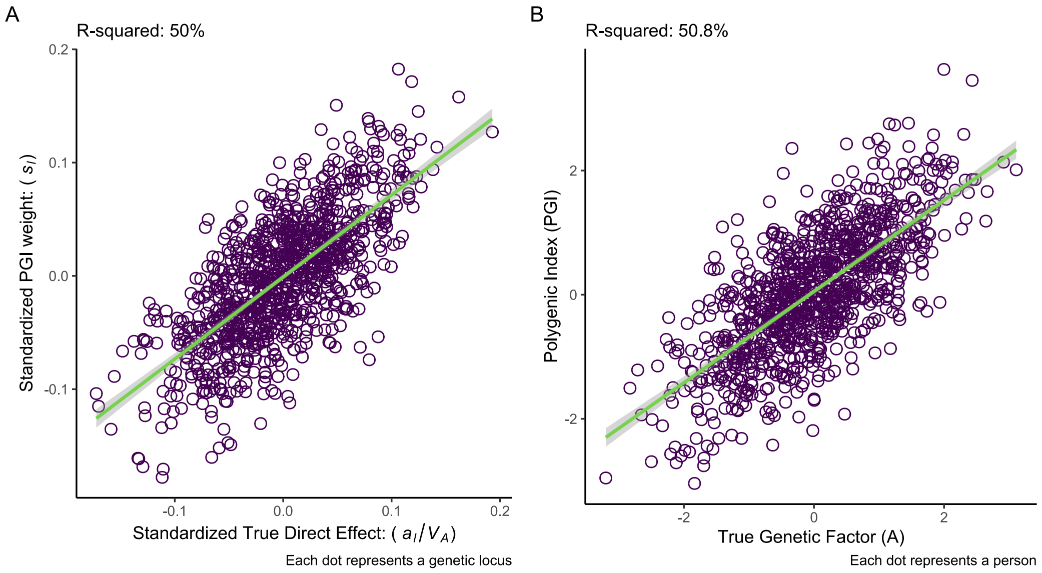

**Supplementary Figure 15***Simulated example of the quality of a polygenic index. With no assortative mating, the correlation between the true genetic effects and the polygenic index weights (Panel A) will be the same as the correlation between individuals' polygenic indices and their true genetic factor (Panel B).*

In Note 5.6, we simulate biased polygenic weights.

#### 4.5 Setting the parameters

The simulations assumed constant partner correlations across generations and constant causal effects of genes and environments. Increasing phenotypic variance thus comes from increased genetic variance and environmental variance. Because we are interested in what the consequences of assortative mating have been (and not what they would be if it started now), we set the parameters with respect to the equilibrium population, and calculated what the starting values in the base population should be to end up at the set parameters. In all simulations, we set the parameters so that the phenotypic variance would end up having unit variance in the equilibrium population, meaning that for example the genetic variance at equilibrium directly corresponds to the heritability.

The user-set parameters were as follows:

- the partner correlation, $\mu$
- the equilibrium additive heritability, $V_A(\infty) = h^2(\infty)$
- the dominance heritability, $V_D = d^2(\infty)$
- the causal effect of the parental environment on the offspring environment, $b$
- the causal effect of the parental phenotype on the offspring environment, $p$
- the generation-specific environmental variance shared by siblings, $V_C$
- the equilibrium genetic signal, $s^2(\infty)$

Because the gene-environment covariance, $V_\omega(\infty)$, and environmental variance, $V_E(\infty)$, are mutually dependent, they were calculated simultaneously using the other set parameters so that the phenotypic variance would end up at unit variance. From this, we could also derive the environmental variance unique to each individual, $V_\epsilon$. Using the set and derived parameters, we calculated what $V_A(0)$, $V_E(0)$, $V_\omega(0)$ and $s^2(0)$ should be in the base population so that they would end up as expected in the equilibrium population (see script at https://osf.io/dgw4r/). These values were then used as input for the simulation.

The script we provide allows the user to set some other properties themselves, such as the sample size and number of loci. In the simulations we present here, we simulated $n_{pairs} = 50{,}000$ (meaning a total population size of 100,000 per generation), each with $n_{loci} = 1000$ loci distributed on $n_{chr} = 10$ chromosomes. The minor allele frequencies were set to $q_l = .50$ for all loci, but the provided script includes an option to have random allele frequencies. We simulated 15 generations of assortment in each run (in addition to the base population and founder population)

### Supplementary Note 5 Simulations: Results

#### 5.1 Genetic transmission only

In the first simulation, we simulated genetic transmission only. That is, the only reason family members resemble each other is because of shared additive genetic factors. We set the partner correlation to $\mu = .50$ and the equilibrium heritability to $h^2(\infty) = 50\%$ (meaning $e^2(\infty) = 50\%$). These are the same values used as examples in the main paper. As expected, the genetic variance increased in the first few generations of assortative mating before stabilizing (Supplementary Figure 16A). In this example, the environmental variance remained constant across generations, meaning that the heritability – the *relative* importance of genetic factors to total phenotypic variance – increased as well (Supplementary Figure 16B).

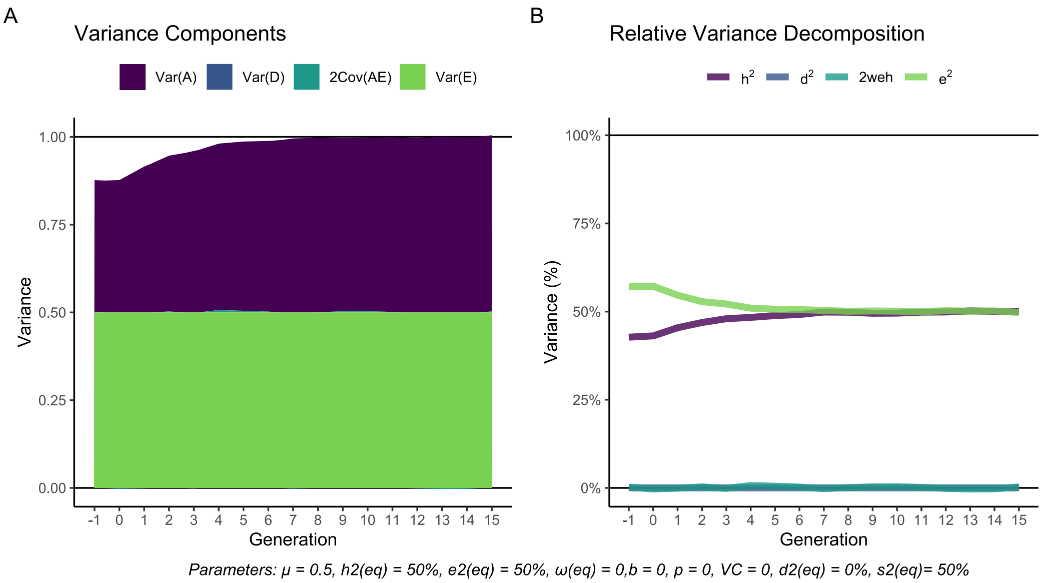

**Supplementary Figure 16** *Simulation of different sources of variance across generations with* $h^2(\infty) = 50\%$ *and a constant partner correlation of* $\mu = .50$ *from generation 0 and onwards.*

Supplementary Figure 17 presents the true genotypic correlations between relatives (purple line) in successive generations of assortative mating. The grey dashed lines are the equilibrium correlations we would expect based on the simulated parameters and on the equations presented in earlier notes. All correlations conform to these expectations. We also see that – as predicted – the correlations between relatives start off at the coefficient of relatedness (black solid lines) and increase in the first few generations of assortment before stabilizing at an equilibrium. In this example, it happens after approximately 6 generations, although we saw in Supplementary Note 2 that this will depend on the heritability and strength of assortment.

The green line is the predicted equilibrium correlation between relatives using Equation (5) from the main paper ($r_{g_k} = \left(\frac{1+\rho_g}{2}\right)^k$). We can see that the true correlation (purple) matches the predicted correlation (green), but only in equilibrium. Before equilibrium, the true correlation is less than Equation (5) would predict. This is just as predicted given the theoretical expectations laid out in Note 2.3.

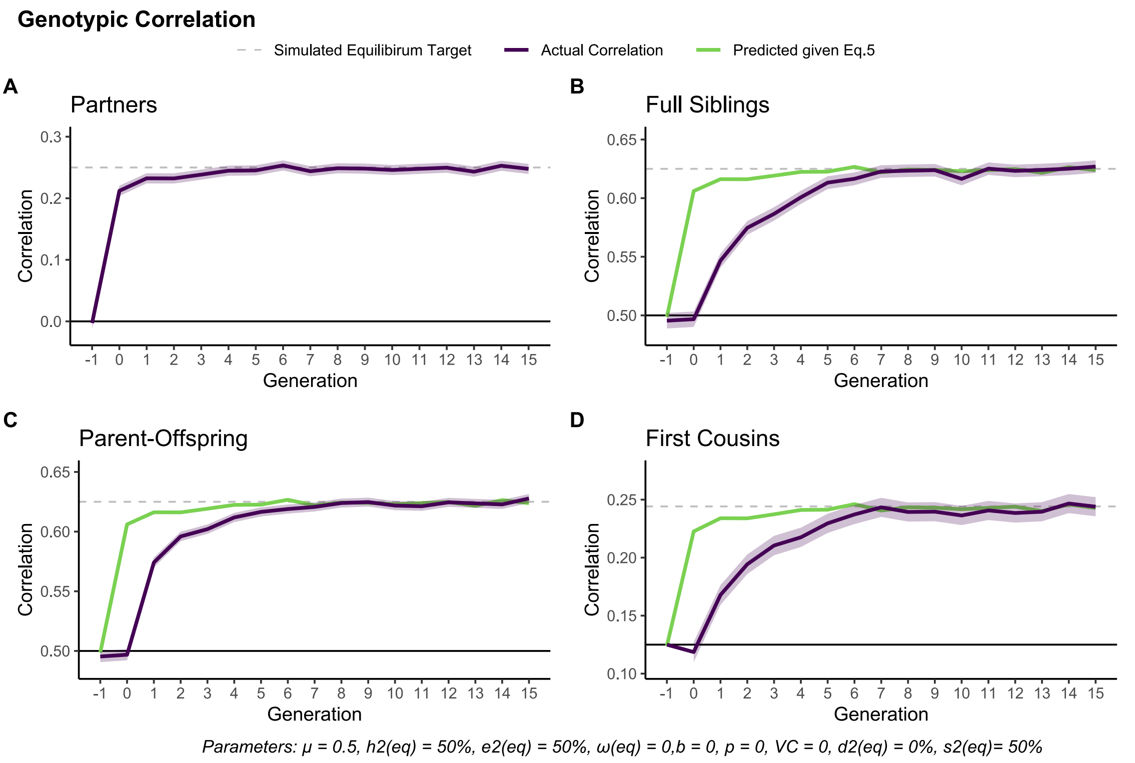

**Supplementary Figure 17** *Simulation of genotypic correlations (purple, with 95% CIs) between family members across generations with* $h^2(\infty) = 50\%$ *and a constant partner correlation of* $\mu = .50$ *from generation 0 and onwards. The parent-offspring correlation is with parents in the preceding generation. The green lines are the predicted equilibrium correlations between relatives given the true genotypic correlation between partners (Panel A).*

#### 5.2 Dominance effects

In the second simulation, we included dominance effects. In Note 1.1, we saw that we should expect dominance effects to be largely unaffected by assortative mating. To illustrate this, we simulated unrealistically high dominance effects. We set the partner correlation to $\mu = .50$ (again), the equilibrium additive heritability to $h^2(\infty) = 40\%$, and the equilibrium dominance heritability to $d^2(\infty) = 40\%$ (meaning $e^2(\infty) = 20\%$). As we can see in Supplementary Figure 18A, the dominance variance remained constant across generations, and was unaffected by assortative mating. Because dominance variance remained constant while the additive genetic variance increased, the relative importance of dominance effects decreased with successive generations of assortative mating (Supplementary Figure 18B).

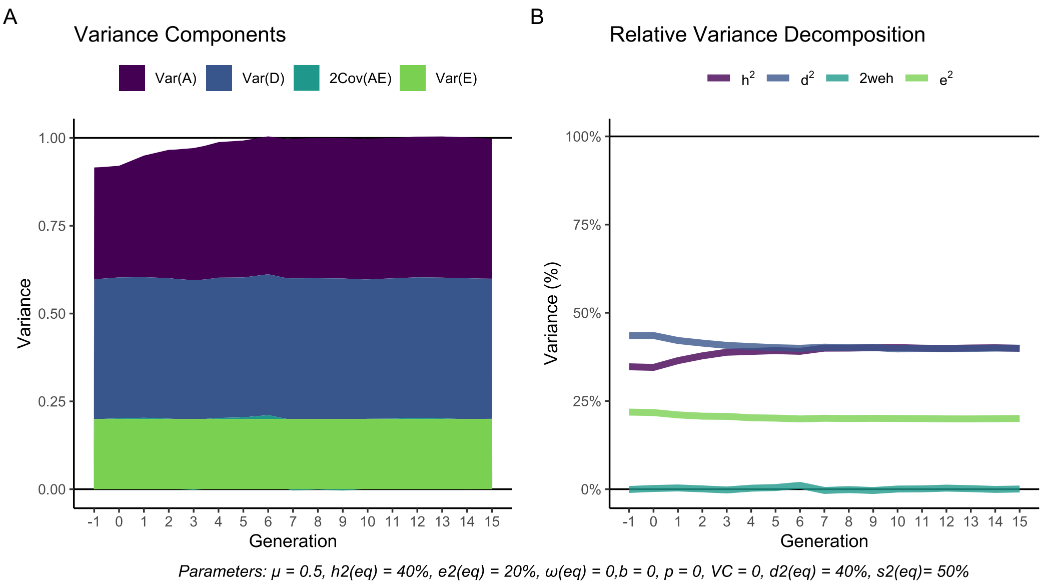

**Supplementary Figure 18** *Simulation of different sources of variance across generations with* $h^2(\infty) = 40\%$, $d^2(\infty) = 40\%$ *and a constant partner correlation of* $\mu = .50$ *from generation 0 and onwards.*

When it comes to the additive genotypic correlations, we predicted in Note 1.1 that they would remain largely unaffected. First-degree relatives should be completely unaffected, and the effect on the genotypic correlation between first cousins should be negligible. As we can see in Supplementary Figure 19, additive genotypic correlations between relatives behave just like when no dominance effects were simulated. The effect on first cousins is too small to detect, even with the unrealistically large dominance effects.

We also expected that the correlation between relatives' dominance effects would remain unaffected by assortative mating. In Supplementary Figure 20, we that this is indeed the case. The dominance correlation between partners (Panel A) was, as expected, the product of the phenotypic correlation and the dominance heritability. The correlation between siblings (Panel B) remained at .25, whereas the correlation between parents and offspring (Panel C) and between first cousin (Panel D) remained at 0.

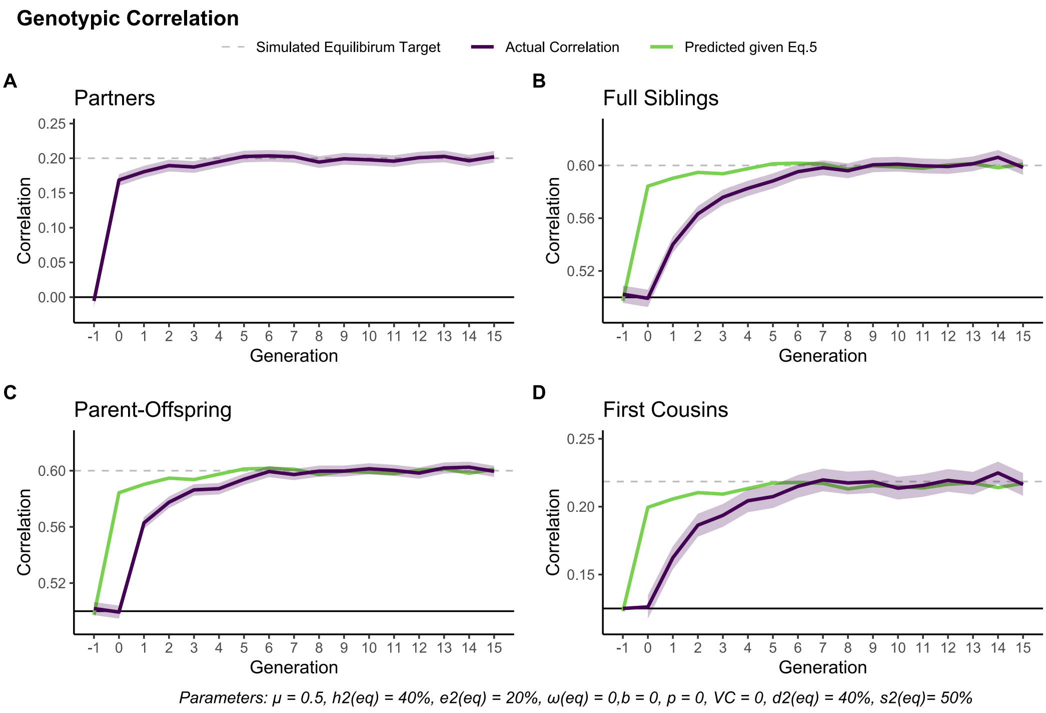

**Supplementary Figure 19** *Simulation of genotypic correlations (purple, with 95% CIs) between family members across generations with* $h^2(\infty) = 40\%$, $d^2(\infty) = 40\%$ *and a constant partner correlation of* $\mu = .50$ *from generation 0 and onwards. The parent-offspring correlation is with parents in the preceding generation. The green lines are the predicted equilibrium correlations between relatives given the true genotypic correlation between partners (Panel A).*

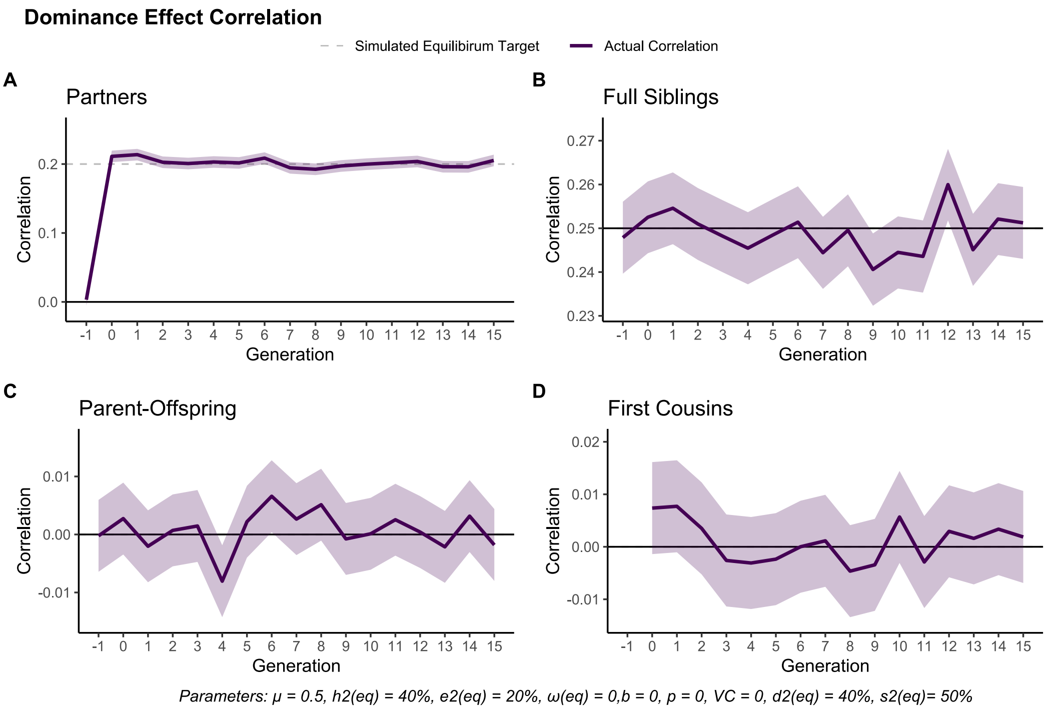

**Supplementary Figure 20** *Simulation of dominance correlations (with 95% CIs) between family members across generations with* $h^2(\infty) = 40\%$, $d^2(\infty) = 40\%$ *and a constant partner correlation of* $\mu = .50$ *from generation 0 and onwards. The parent-offspring correlation is with parents in the preceding generation.*

#### 5.3 Environmental effects shared by siblings

In the third simulation, we replaced the dominance effects with environmental effects shared by siblings. As described in Note 1.1, this should have similar effects as dominance effects, only potentially more noticeable. We set the partner correlation to $\mu = .50$ (again) and the equilibrium heritability to $h^2(\infty) = 40\%$ (meaning $e^2(\infty) = 60\%$). We set the parameters so that siblings would share 2/3 of the environmental variance ($V_C = .4$, equivalent to 40% shared environmental variance). This is the same as the example values used on page 6. Note that this was only shared within each generation and did not contribute to parent-offspring similarity or cousin similarity.

As we can see in Supplementary Figure 21, genetic similarity in the nuclear family is unaffected by environmental effects shared by siblings. For first cousins, however, we see that the true genotypic correlation (the purple line in Panel D) at equilibrium was slightly above the correlation one would predict by using Equation (5). After 15 generations, the true genotypic correlation was .230 (95% CIs: .221, .238), whereas using Equation (5) would predict $\left(\frac{1+\rho_g}{2}\right)^3 = \left(\frac{1+.206}{2}\right)^3 = .219$. These values match the theoretical expectation described on page 6.

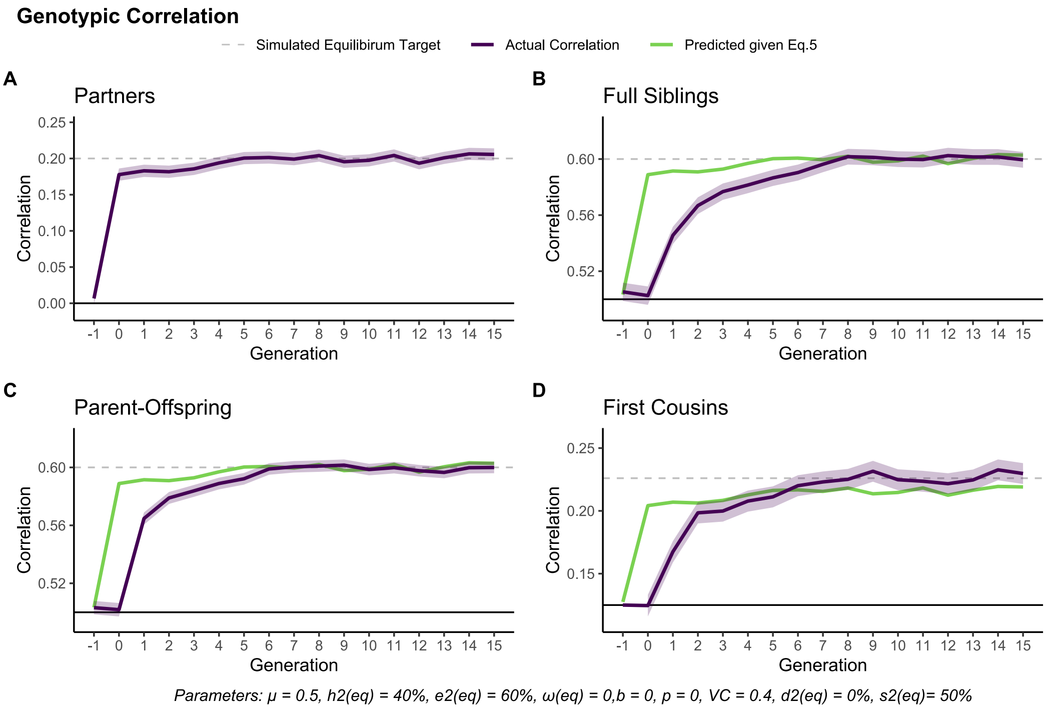

**Supplementary Figure 21** *Simulation of genotypic correlations (purple, with 95% CIs) between family members across generations with $h^2(\infty) = 40\%$, $V_C = .4$ and a constant partner correlation of $\mu = .50$ from generation 0 and onwards. The parent-offspring correlation is with parents in the preceding generation. The green lines are the predicted equilibrium correlations between relatives given the true genotypic correlation between partners (Panel A).*

#### 5.4 Environmental transmission and gene-environment correlations

In the fourth simulation, we simulated intergenerational environmental transmission. As with the other simulations, we used the example values used when illustrating the theoretical expectations (see page 11). That is, we set the partner correlation to $\mu = .50$, the equilibrium heritability to $h^2(\infty) = 40\%$, the effect of the parental environment to $b = .20$, and finally the effect of the parental phenotype to $p = .10$. For the phenotypic variance to be 1 at equilibrium, the environmental effects at equilibrium would have to be $e^2 = 34\%$ and the gene-environment correlation would have to be $\omega = .35$.

As expected, both genetic variance, environmental variance, and the gene-environment correlation increased with successive generations of assortative mating (Supplementary Figure 22A). For these parameters, the variance attributable to gene-environment covariance (coloured teal) increased relatively faster than the other variance terms, which in turn means that the heritability (purple line in Supplementary Figure 22B) decreased with successive generations of assortative mating until it reached equilibrium.

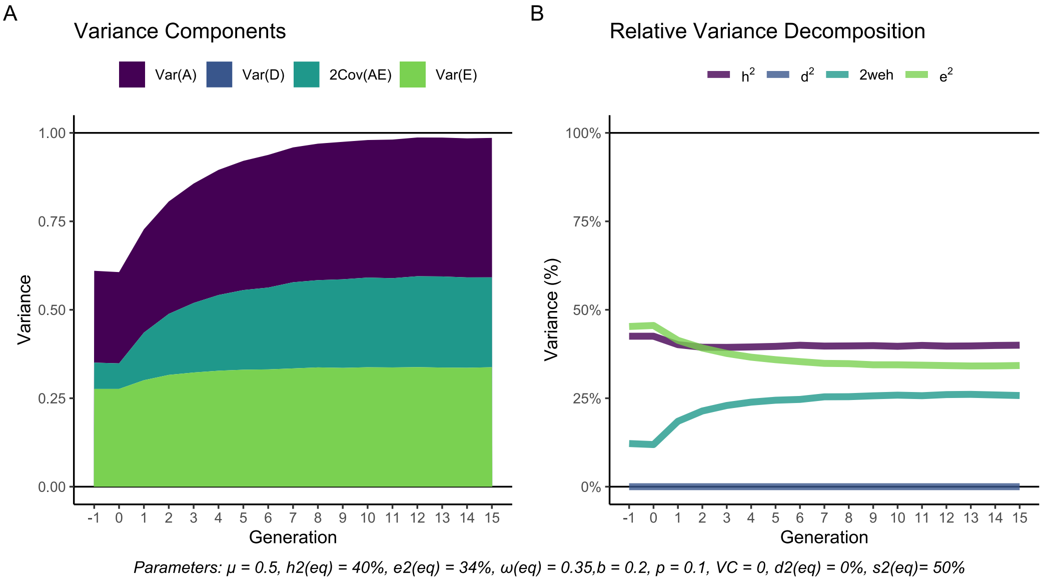

**Supplementary Figure 22** *Simulation of different sources of variance across generations with* $h^2(\infty) = 40\%$, $b = .20$, $p = .10$ *and a constant partner correlation of* $\mu = .50$ *from generation 0 and onwards.*

The genotypic correlations between family members became, as expected, greatly increased in the presence of gene-environment correlations (Supplementary Figure 23). First and foremost, we see that the true genotypic correlation between partners was much larger than in the previous two simulations, despite the same heritability (Panel A). We also see that the partner correlation took a few generations to reach the correlation predicted by the simulated parameters (grey dashed line). This is for the same reason as described on page 16, namely that the genotypic partner correlation is a function of the correlation between the genotype and phenotype ($h + \omega e$), which in turn is a function of the genotypic partner correlation in preceding generations. This feedback-loop increases the genotypic partner correlation further and leads to a few more generations in disequilibrium.

We also see that the genotypic correlation between first-degree relatives (Panel B and C) was greatly increased. However, we also see that Equation (5) (the green line) correctly predicts the correlation at equilibrium. For first cousins (Panel D), on the other hand, gene-environment correlations resulted in a true genotypic correlation above what Equation (5) would predict.

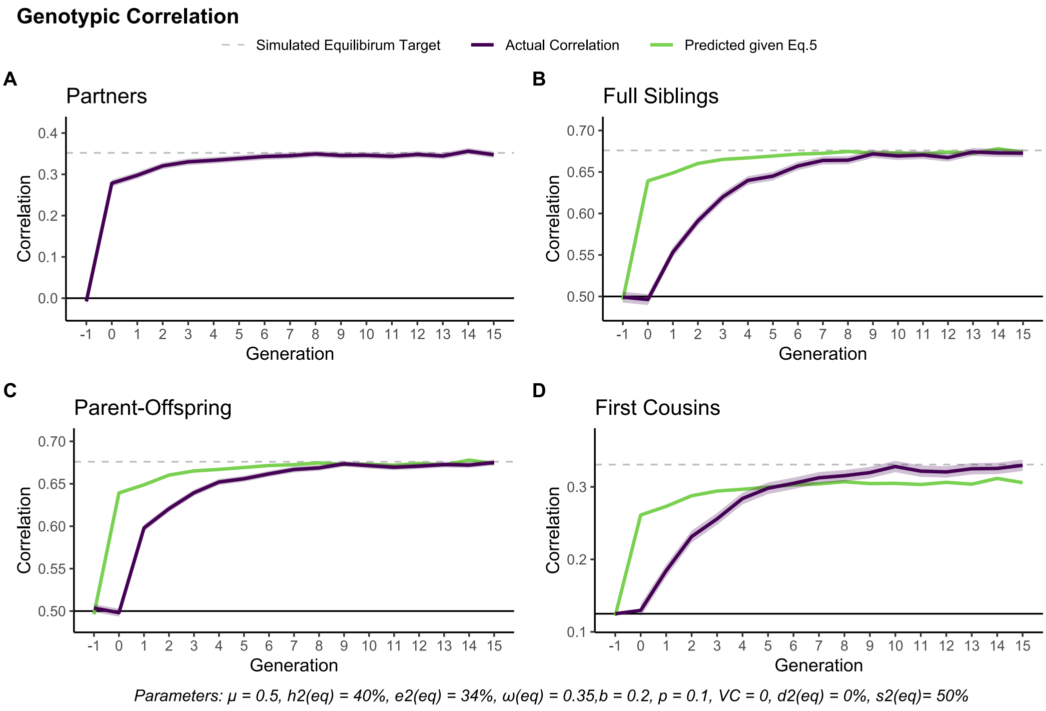

**Supplementary Figure 23** *Simulation of genotypic correlations (purple, with 95% CIs) between family members across generations with* $h^2(\infty) = 40\%$, $b = .20$, $p = .10$ *and a constant partner correlation of* $\mu = .50$ *from generation 0 and onwards. The parent-offspring correlation is with parents in the preceding generation. The green lines are the predicted equilibrium correlations between relatives given the true genotypic correlation between partners (Panel A).*

#### 5.5 Polygenic indices

In all four simulations described up to now, we also simulated polygenic indices with a target genetic signal in the equilibrium population set to $s^2(\infty) = 50\%$. Here we present the results for the polygenic indices for two of these simulations: The first (genetic transmission only) and the fourth (genetic and environmental transmission). In the first simulation, the parameters would imply a genetic signal in the base population of $s^2(0) = 43\%$, whereas in the fourth simulation, the parameters would imply$s^2(0) = 39\%$. In Supplementary Figure 24, we see the squared correlation between individuals' polygenic indices and their true genetic factors (i.e., the genetic signal). As expected, we see that the genetic signal rises in successive generations of assortative mating until it reaches the equilibrium.

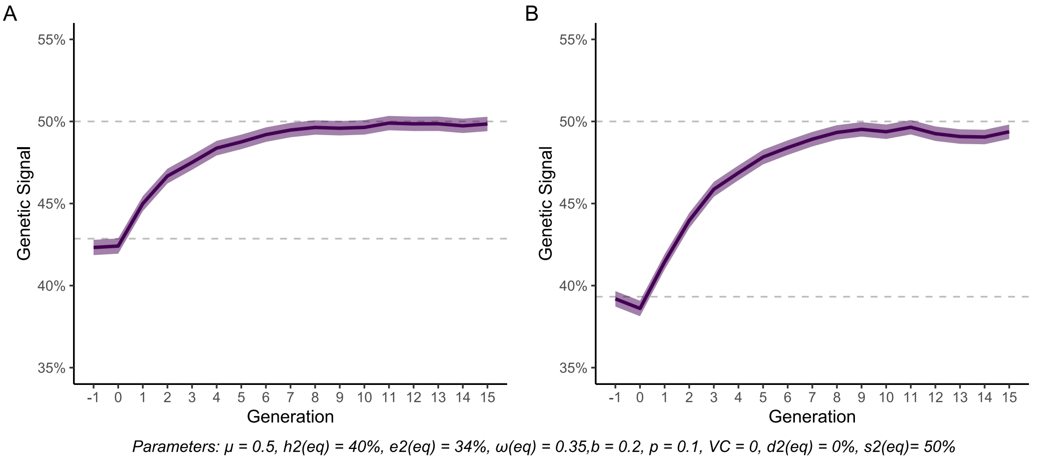

**Supplementary Figure 24** *Genetic signal across successive generations of assortative mating ($\mu = .50$). **A**: Genetic transmission only,* $h^2(\infty) = 50\%$ *and* $\omega(\infty) = 0$*; **B**: Genetic and environmental transmission,* $h^2(\infty) = 40\%$ *and* $\omega(\infty) = .35$.

When it comes to the correlations between family members' polygenic indices, we see that they are attenuated compared to the true genotypic correlations (Genetic transmission only: Supplementary Figure 25, *cf. Supplementary Figure 17*; Genetic and environmental transmission: Supplementary Figure 26, *cf. Supplementary Figure 23*). However, the correlations still conform to the equations laid out it earlier, namely that the polygenic index correlations are the true genotypic correlations weighted by the genetic signal. Within the nuclear family, we see that the relationship between the correlation between partners and the correlations between relatives are the same as for polygenic indices as for the true genetic factor. For first cousins, the true polygenic index correlation (purple line) is higher than expected by Equation (5) ($r_{g_3} = \left(\frac{1+\rho_g}{2}\right)^3$, green line). For a situation with genetic transmission only (Supplementary Figure 25D), this discrepancy is barely noticeable. However, it is very noticeable for a situation with both genetic and environmental transmission (Supplementary Figure 26D). This is because the accuracy of Equation (5) is off both because of poor genetic signal *and* gene-environment correlations.

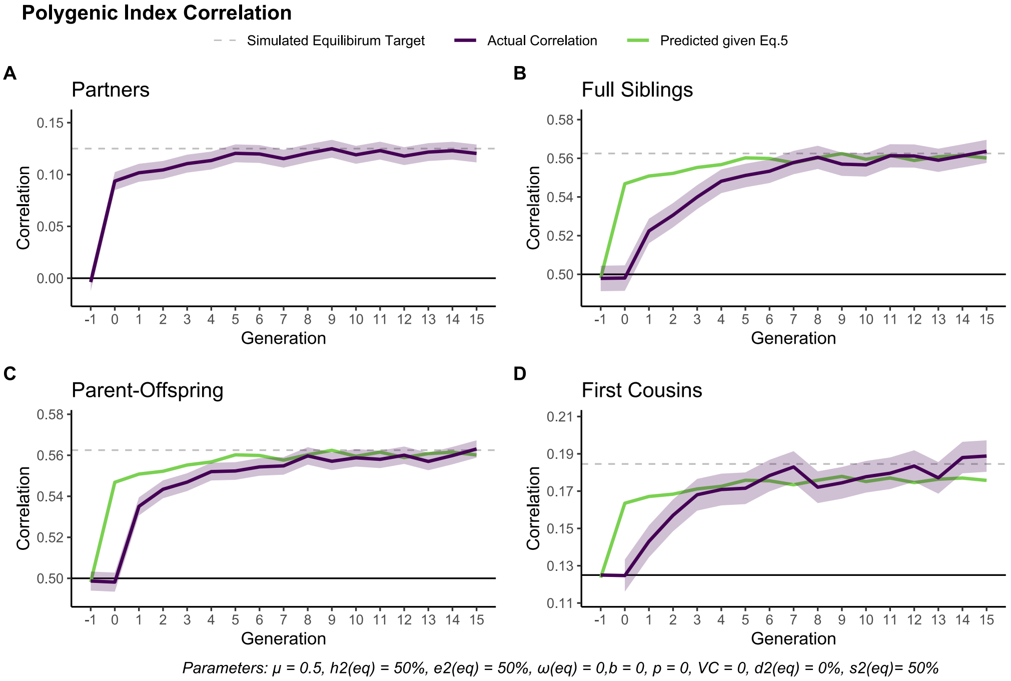

**Supplementary Figure 25** *Simulation of polygenic index correlations (purple, with 95% CIs) between family members across generations with* $h^2(\infty) = 50\%$, $s^2(\infty) = 50\%$, *and a constant partner correlation of* $\mu = .50$ *from generation 0 and onwards. The parent-offspring correlation is with parents in the preceding generation. The green lines are the predicted equilibrium polygenic index correlations between relatives given the polygenic index correlation between partners (Panel A).*

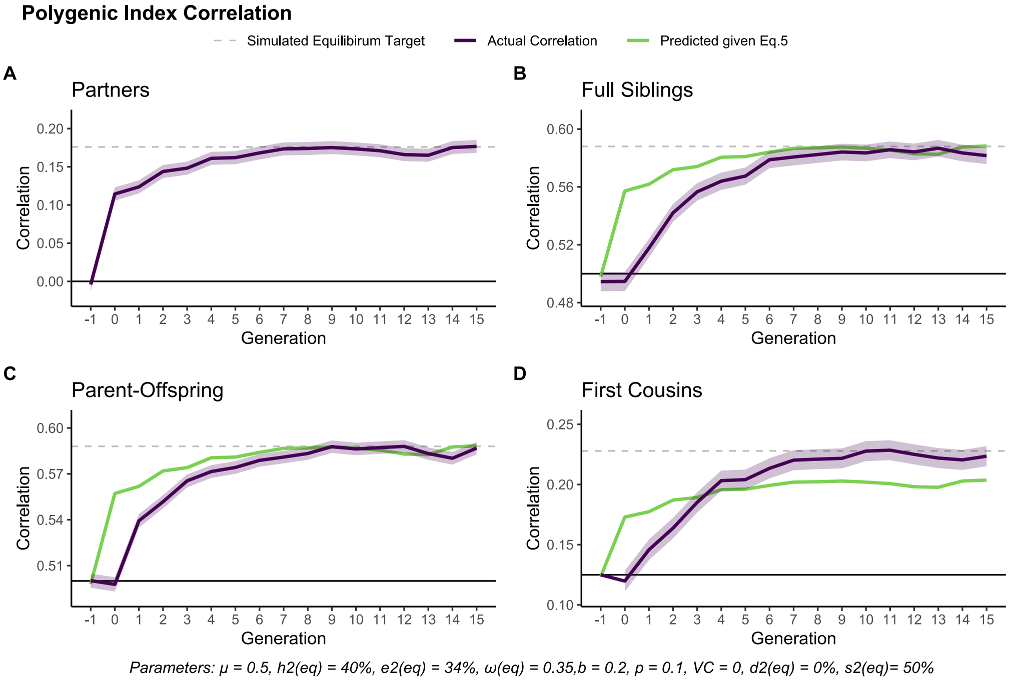

**Supplementary Figure 26** *Simulation of polygenic index correlations (purple, with 95% CIs) between family members across generations with* $h^2(\infty) = 40\%$, $\omega(\infty) = .35$, $s^2(\infty) = 50\%$, *and a constant partner correlation of* $\mu = .50$ *from generation 0 and onwards. The parent-offspring correlation is with parents in the preceding generation. The green lines are the predicted equilibrium polygenic index correlations between relatives given the polygenic index correlation between partners (Panel A).*

#### 5.6 Polygenic indices based on biased weights

In the simulations above, the weights ($s_l$) used to calculate polygenic indices were noisy, but unbiased. However, under assortative mating, weights obtained from genome-wide association studies (GWAS) will most likely be biased. This occurs because assortative mating induces linkage disequilibrium between trait-associated loci, meaning that each locus is associated with all the other trait-associated loci. This increases the association between each locus and the phenotype, leading to upwardly biased weights, and in turn, biased polygenic indices[21]. A key concern is that this also biases correlations between relatives. If this bias differentially affects close and distant relatives, or if it affects the relationship between the partner correlation and the correlation between relatives, then our expectations laid out in Supplementary Note 3 will be inaccurate and our results may be misleading.

##### ***Simulating biased GWAS analyses***

To make sure this was not the case, we needed to simulate biased polygenic index weights. To accomplish this, we ran a new simulation using the same parameters as in the first simulation (i.e., a partner correlation at $\mu = .50$ and equilibrium heritability of $h^2_\infty = 50\%$). Instead of merely adding noise to the true genetic effects, we calculated polygenic index weights by doing a GWAS on a subsample of 5000 individuals. More specifically, we let the new weights be the regression slopes of the phenotype on each locus. To prevent individuals in the discovery sample also being included in the analyses below, we simulated additional individuals in the generations where the weights were estimated. For comparisons sake, we estimated new weights three times during the simulation: in the base population (i.e., random mating), after 2 generations of assortment (i.e., disequilibrium), and finally after 15 generations of assortment (i.e., equilibrium). The new weights were used to calculate polygenic indices from that generation onward, using equation (S4.8a).

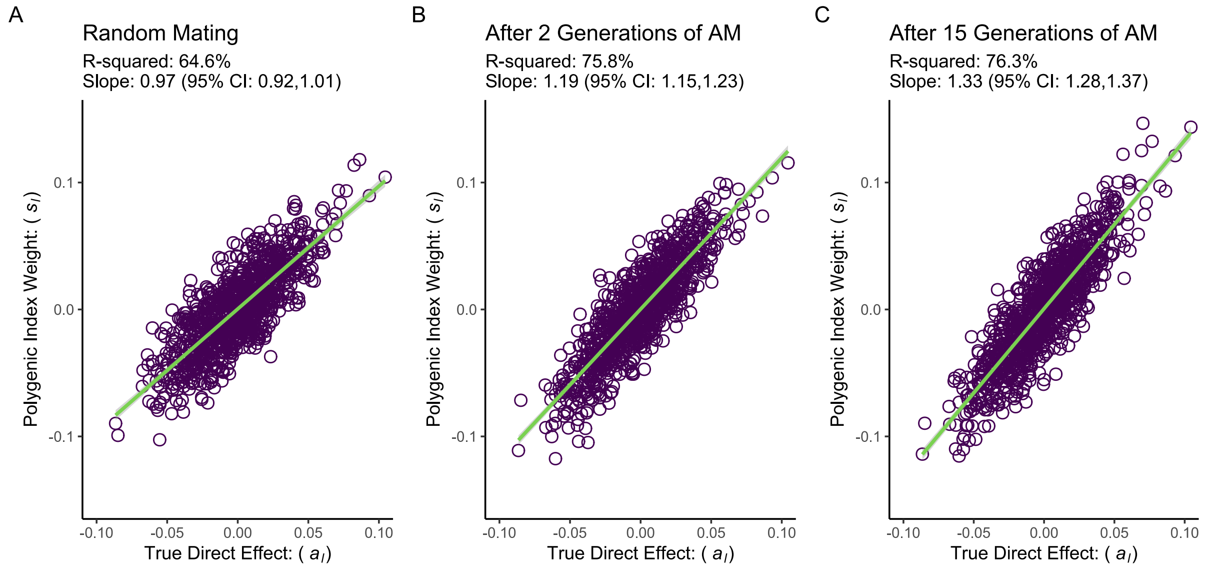

**Supplementary Figure 27** *Quality of polygenic index weights estimated by regressing a simulated phenotype on each locus. The figure shows regressions of estimated polygenic index weights on true genetic effects after 0 (A), 2 (B), and 15 (C) generations of assortative mating with a partner correlation of $\mu = .50$ and an equilibrium heritability of $h^2_\infty = 50\%$. Unbiased weights should have a unit slope, while a slope >1 indicates overestimated weights.*

Supplementary Figure 27 illustrates the results of regressing the estimated polygenic index weights ($s_l$) on the true genetic effects ($a_l$). In the random mating population (Panel A), we see that the regression slope is not significantly different from a unit slope, which means that the weights are unbiased. This is not the case under assortative mating. For example, at equilibrium (Panel C), the slope is 1.33, indicating that the polygenic index weights substantially overestimate the genetic effects. Note also that the shared variance ($R^2$) between the true effects and the weights increases with assortative mating. This occurs because the sampling error is the same across the three analyses, which counterintuitively means that the biased weights are more highly correlated with the true effects than the unbiased weights. This is further exacerbated in the genetic signal: In a random mating population, using the unbiased weights, we unsurprisingly found that the genetic signal equalled the shared variance between the true effects and the weights, $s^2 = 64.6\%$ $(95\%\ CIs: 64.2\%, 64.9\%)$. However, after two generations of assortative mating (using the new weights estimated in that generation), the genetic signal had increased to $s^2 = 78.8\%$ $(78.6\%, 79.1\%)$, and after 15 generations (again using the new weights from that generation), the genetic signal had reached $s^2 = 81.2\%$ $(81.0\%, 83.4\%)$.

##### *Effects on correlations between relatives at equilibrium*

To investigate whether this bias affects our expectations about similarity among family members, we first investigated correlations at equilibrium. We did this by simulating a 16th generation (so that we would have parents and offspring with polygenic indices using the same weights). According to Supplementary Note 3 (page 23), we would expect polygenic index correlations between partners to follow equation (S3.1), while polygenic index correlations between $k^{\text{th}}$-degree relatives should follow equation (S3.2). Key to both equations is that the true genotypic correlation should be attenuated by the genetic signal (which in generation 16 was practically the same as in the generation before: $s^2 = 81.048\%$). We found that correlations between polygenic indices conformed to the expectations laid out earlier:

- The true genotypic correlation between partners was $\rho_g = .251$ $(.242, .259)$, meaning we would expect a polygenic index correlation of $.203$. This matches the observed polygenic index correlation: $\rho_{pgi} = .208$ $(.199, .216)$.
- For first-degree relatives, there are two ways we could calculate the expected polygenic index correlation: The first is using the polygenic index correlation between partners, meaning $\frac{1+.203}{2} = .604$. The other is using the true genotypic correlation ($r_g = .624$ for both types of relatives) and the genetic signal, meaning the expected polygenic index correlation is $.601$. Both of these expectations match what we observe: $r_{pgi} = .603$ $(.599, .607)$ for parents and offspring, and $r_{pgi} = .602$ $(.596, 608)$ for full siblings.
- Finally, the true genotypic correlation between first cousins was $r_{g_3} = .242$ $(.234, .250)$, meaning we would expect a polygenic index correlation of $s^2 r_{g_3} + (1 - s^2)\left(\frac{1}{2}\right)^3 = .220$. This also matches the observed polygenic index correlation $r_{pgi} = .219$ $(.211, .227)$.

##### *Effects on measures of disequilibrium*

To investigate whether expectations also hold during disequilibrium, we calculated intergenerational variance ratios ($Q$) and observed-versus-expected increases in the parent-offspring correlation ($U$) on the simulated data, both for the true genetic factors and for the polygenic indices. All results were as expected given the expectations laid out earlier:

- <u>Random mating population</u>: Here, as expected, both the true genetic variance and the polygenic index variance were constant across generations, with the variance ratios being $Q_A = 0.997$ $(0.988, 1.006)$ and $Q_{pgi} = 1.000$ $(0.991, 1.009)$, respectively. $U$ will involve dividing by near-zero in a random mating population and is therefore not informative here.
- <u>Disequilibrium population:</u> When comparing variance after 2 and 3 generations of assortative mating (using the polygenic index weights estimated after 2 generations), we found that true genetic variance was larger in the offspring generation, $Q_A = 1.033$ $(1.026, 1.040)$. For polygenic index variance, the variance ratio was smaller, but still indicated greater variance in the offspring generation, $Q_{pgi} = 1.022$ $(1.015, 1.030)$. Given the genetic signal reported above, this value is as expected given equation (S3.13). When it comes to $U$, we found similar values for both the true genotypic correlation and the polygenic index: $U_A = 91.4\%$ $(89.7\%, 93.1\%)$ and $U_{pgi} = 92.3\%$ $(90.7\%,\ 95.2\%)$, respectively. They were broadly as expected after two generations of assortative mating, albeit a bit on the high side (see Note 2.3).

- Equilibrium population: When comparing variance after 15 and 16 generations of assortative mating (using weights estimated after 15 generations), we again found that both the true genetic variance and the polygenic index variance were constant across generations, with the variance ratios being $Q_A = 1.001\ (0.995, 1.008)$ and $Q_{pgi} = 0.996\ (0.989, 1.003)$, respectively. Furthermore, $U$ was at 100% for both the true genotypic correlation ($U_A = 100.1\%\ (99.5\%, 100.8\%)$) and the polygenic index corelation ($U_{pgi} = 99.6\%\ (98.9\%, 100.3\%)$).

In other words, the expectations laid out in Supplementary Note 3 still appear to hold, even for biased polygenic indices. Correlations between close and distant relatives appear not be differentially biased, nor does the bias appear to affect the relationship between the partner correlation and the correlation between relatives. Likewise, conclusions about disequilibrium does not appear to be very affected by biased polygenic indices.

#### Supplementary Note 6 Empirically testing intergenerational equilibrium

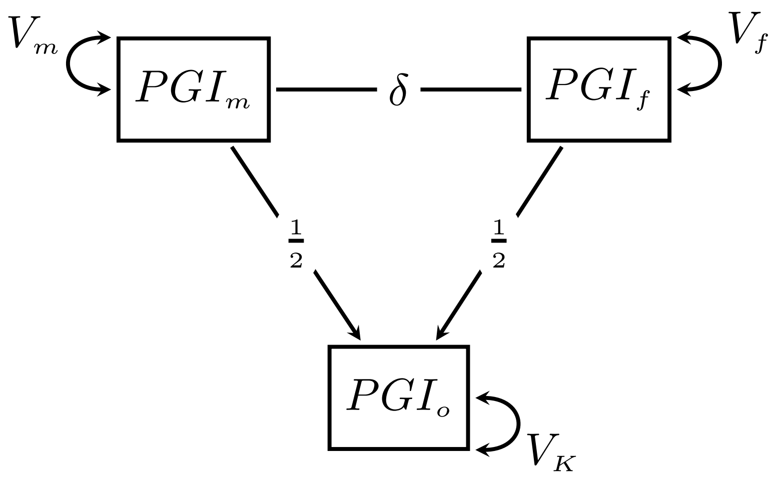

**Supplementary Figure 28** *Structural equation model used to test intergenerational equilibrium.*

Whether a trait is consistent with equilibrium can be quickly gauged by seeing if the confidence intervals for the correlations in Fig. 3 overlaps with the expected correlations (the black crosses). To perform a more formal test of equilibrium, we fitted structural equation models (Supplementary Figure 28) to mother-father-offspring trios for each trait and used a log-likelihood test to check if constraining the model to equilibrium (i.e., equal variance across generations) resulted in significantly poorer fit. This approach also allowed us to compute $Q_{pgi}$ and $U_{pgi}$ with confidence intervals, which offer two alternative ways to check for equilibrium: First, whether the observed parent-offspring correlation was smaller than expected given the partner correlation (i.e., if $U_{pgi} \neq 1$, ref. Equation (S3.18)). Second, whether the variance in the offspring generation was greater than in the parental generation (i.e., $Q_{pgi} \neq 1$, ref. Equation (S3.8)). Note that these are not independent tests, but rather transformations of each other that highlight different aspects of disequilibrium. The models were instantiated in OpenMx[23] in R[24].

To avoid putting needless constraints on the model, the maternal and paternal variance ($V_m$ and $V_f$, respectively) were allowed to differ. The partner covariance is therefore $V_m \delta V_p$, whereas the parent-offspring covariances are $\frac{V_m+V_m\delta V_f}{2}$ and $\frac{V_f+V_m\delta V_f}{2}$, respectively (For $U_{pgi}$ and $Q_{pgi}$, we used the combined variance, see below). The offspring variance is slightly more complicated: $V_K + \frac{V_m}{4} + \frac{V_f}{4} + \frac{V_m\delta V_f}{2}$. Note that these equations are the same as those presented in Supplementary Note 2, only adapted for varying maternal and paternal variance.

Both the parent-offspring correlations and the intergenerational variance ratio will depend on the recombination variance ($V_K$). The recombination variance will be half the genetic variance in the parent generation in the absence of assortative mating (Equation S2.2). For a trait at equilibrium, $V_K$ can be found by rearranging Equation S2.9b and substituting it into Equation (S2.2). To estimate how much the recombination variance deviated from what would be expected at equilibrium, we added a new freely estimated parameter ($x$) to this equation:

$$V_K = \frac{\sqrt{V_f V_m} + V_m \delta V_f + x}{2} \qquad (S6.1)$$

This parameter allows the variance in the offspring generation to differ from the parent generation. If $x$ is significantly different from zero, then the trait is not consistent with equilibrium. This can be tested by constraining $x$ to 0 and seeing if it results in significantly worse fit (see Supplementary Tables 1 through 16 below).

We also defined $Q_{pgi}$ and $U_{pgi}$ in the model, which is how we obtained confidence intervals for these parameters. If $x > 0$, then $Q_{pgi} > 1$ and $U_{pgi} < 1$. To define $U_{pgi}$, we also needed to define the partner correlation ($\rho_{pgi}$) and the pooled parent-offspring correlation ($r_{pgi}$). Note again that these equations are the same as those presented earlier, only adapted for varying maternal and paternal variance:

$$\rho_{pgi} = \sqrt{V_m V_f} \times \delta \qquad (S6.2)$$

$$r_{pgi} = \frac{\left(\sqrt{V_m V_f} + V_m \delta V_p\right)/2}{\sqrt{\sqrt{V_m V_f} \times Var(PGI_o)}} \qquad (S6.3)$$

$$U_{pgi} = \frac{r_{pgi} - 0.5}{\left(1 + \rho_{pgi}\right)/2 - 0.5} \qquad (S6.4)$$

$$Q_{pgi} = \frac{Var(PGI_o)}{\sqrt{V_m V_f}} \qquad (S6.5)$$

We report $U_{pgi}$ and $Q_{pgi}$ for traits with significant partner correlations in Fig. 4 in the main paper.

We also used this modelling approach as an alternative test for significant partner similarity (i.e., assortative mating) by testing whether constraining $\delta$ to 0 resulting in significantly worse fit (after constraining the model to equilibrium). None of the models contradicted the results reported in the main paper. We further compared a model where we constrained $V_f$ to equal $V_m$ (i.e., sex invariant variance). Significant differences would indicate that men and women have different selection bias into the sample (or into parenthood). We found this to be the case for chronotype ($p$ = .007), drinking (.020), IQ (<.001), and EA (cognitive) (.012). (This should not affect the equilibrium tests but may be interesting avenues for future research).

**Supplementary Table 1: Model Fit Statistics (and Likelihood Ratio Tests) for ADHD**

| Base | Comparison | *k* | -2LL | df | *Δ* 2LL | *Δ* df | *p* |
|---|---|---|---|---|---|---|---|
| Full | | 4 | 521,217.1 | 195,078 | | | |
| Full | Equilibirium | 3 | 521,217.2 | 195,079 | 0.091 | 1 | 0.7623 |
| Equilibirium | No_Assortment | 2 | 521,217.3 | 195,080 | 0.046 | 1 | 0.8293 |
| No_Assortment | Sex_Invariant | 1 | 521,218.5 | 195,081 | 1.200 | 1 | 0.2734 |

**Supplementary Table 2: Model Fit Statistics (and Likelihood Ratio Tests) for Bipolar Disorder**

| Base | Comparison | *k* | -2LL | df | *Δ* 2LL | *Δ* df | *p* |
|---|---|---|---|---|---|---|---|
| Full | | 4 | 521,403.9 | 195,082 | | | |
| Full | Equilibirium | 3 | 521,410.7 | 195,083 | 6.811 | 1 | 0.0091 |
| Equilibirium | No_Assortment | 2 | 521,412.2 | 195,084 | 1.414 | 1 | 0.2343 |
| No_Assortment | Sex_Invariant | 1 | 521,412.6 | 195,085 | 0.480 | 1 | 0.4886 |

| Supplementary Table 3: Model Fit Statistics (and Likelihood Ratio Tests) for BMI | | | | | | | |
|---|---|---|---|---|---|---|---|
| **Base** | **Comparison** | ***k*** | **-2LL** | **df** | **Δ 2LL** | **Δ df** | ***p*** |
| Full | | 4 | 520,633.4 | 195,116 | | | |
| Full | Equilibirium | 3 | 520,657.8 | 195,117 | 24.373 | 1 | 0.0000 |
| Equilibirium | No_Assortment | 2 | 520,712.2 | 195,118 | 54.438 | 1 | 0.0000 |
| No_Assortment | Sex_Invariant | 1 | 520,713.1 | 195,119 | 0.917 | 1 | 0.3382 |

| Supplementary Table 4: Model Fit Statistics (and Likelihood Ratio Tests) for Chronotype | | | | | | | |
|---|---|---|---|---|---|---|---|
| **Base** | **Comparison** | ***k*** | **-2LL** | **df** | **Δ 2LL** | **Δ df** | ***p*** |
| Full | | 4 | 519,615.3 | 195,073 | | | |
| Full | Equilibirium | 3 | 519,625.5 | 195,074 | 10.150 | 1 | 0.0014 |
| Equilibirium | No_Assortment | 2 | 519,741.7 | 195,075 | 116.219 | 1 | 0.0000 |
| No_Assortment | Sex_Invariant | 1 | 519,749.0 | 195,076 | 7.256 | 1 | 0.0071 |

| Supplementary Table 5: Model Fit Statistics (and Likelihood Ratio Tests) for Smoking | | | | | | | |
|---|---|---|---|---|---|---|---|
| **Base** | **Comparison** | ***k*** | **-2LL** | **df** | **Δ 2LL** | **Δ df** | ***p*** |
| Full | | 4 | 520,798.9 | 195,157 | | | |
| Full | Equilibirium | 3 | 520,799.1 | 195,158 | 0.192 | 1 | 0.6615 |
| Equilibirium | No_Assortment | 2 | 520,810.0 | 195,159 | 10.978 | 1 | 0.0009 |
| No_Assortment | Sex_Invariant | 1 | 520,812.3 | 195,160 | 2.308 | 1 | 0.1287 |

| Supplementary Table 6: Model Fit Statistics (and Likelihood Ratio Tests) for EA (Cognitive) | | | | | | | |
|---|---|---|---|---|---|---|---|
| **Base** | **Comparison** | ***k*** | **-2LL** | **df** | **Δ 2LL** | **Δ df** | ***p*** |
| Full | | 4 | 519,757.9 | 195,097 | | | |
| Full | Equilibirium | 3 | 519,767.1 | 195,098 | 9.164 | 1 | 0.0025 |
| Equilibirium | No_Assortment | 2 | 519,878.6 | 195,099 | 111.506 | 1 | 0.0000 |
| No_Assortment | Sex_Invariant | 1 | 519,884.9 | 195,100 | 6.353 | 1 | 0.0117 |

| Supplementary Table 7: Model Fit Statistics (and Likelihood Ratio Tests) for Cross-Psychiatric | | | | | | | |
|---|---|---|---|---|---|---|---|
| **Base** | **Comparison** | ***k*** | **-2LL** | **df** | **Δ 2LL** | **Δ df** | ***p*** |
| Full | | 4 | 522,245.9 | 195,094 | | | |
| Full | Equilibirium | 3 | 522,248.9 | 195,095 | 2.972 | 1 | 0.0847 |
| Equilibirium | No_Assortment | 2 | 522,249.1 | 195,096 | 0.270 | 1 | 0.6032 |
| No_Assortment | Sex_Invariant | 1 | 522,250.3 | 195,097 | 1.147 | 1 | 0.2842 |

| Supplementary Table 8: Model Fit Statistics (and Likelihood Ratio Tests) for Drinking | | | | | | | |
|---|---|---|---|---|---|---|---|
| **Base** | **Comparison** | ***k*** | **-2LL** | **df** | **Δ 2LL** | **Δ df** | ***p*** |
| Full | | 4 | 521,337.6 | 195,121 | | | |
| Full | Equilibirium | 3 | 521,338.4 | 195,122 | 0.762 | 1 | 0.3826 |
| Equilibirium | No_Assortment | 2 | 521,345.9 | 195,123 | 7.527 | 1 | 0.0061 |
| No_Assortment | Sex_Invariant | 1 | 521,351.3 | 195,124 | 5.397 | 1 | 0.0202 |

| Supplementary Table 9: Model Fit Statistics (and Likelihood Ratio Tests) for EA | | | | | | | |
|---|---|---|---|---|---|---|---|
| **Base** | **Comparison** | ***k*** | **-2LL** | **df** | **Δ 2LL** | **Δ df** | ***p*** |
| Full | | 4 | 514,606.1 | 195,152 | | | |
| Full | Equilibirium | 3 | 514,642.8 | 195,153 | 36.769 | 1 | 0.0000 |
| Equilibirium | No_Assortment | 2 | 515,755.7 | 195,154 | 1,112.809 | 1 | 0.0000 |
| No_Assortment | Sex_Invariant | 1 | 515,758.5 | 195,155 | 2.801 | 1 | 0.0942 |

**Supplementary Table 10: Model Fit Statistics (and Likelihood Ratio Tests) for Height**

| Base | Comparison | *k* | -2LL | df | *Δ* 2LL | *Δ* df | *p* |
|---|---|---|---|---|---|---|---|
| Full | | 4 | 517,944.8 | 195,053 | | | |
| Full | Equilibirium | 3 | 517,945.5 | 195,054 | 0.691 | 1 | 0.4060 |
| Equilibirium | No_Assortment | 2 | 518,295.6 | 195,055 | 350.158 | 1 | 0.0000 |
| No_Assortment | Sex_Invariant | 1 | 518,296.3 | 195,056 | 0.710 | 1 | 0.3994 |

**Supplementary Table 11: Model Fit Statistics (and Likelihood Ratio Tests) for IQ**

| Base | Comparison | *k* | -2LL | df | *Δ* 2LL | *Δ* df | *p* |
|---|---|---|---|---|---|---|---|
| Full | | 4 | 519,378.7 | 195,095 | | | |
| Full | Equilibirium | 3 | 519,385.0 | 195,096 | 6.223 | 1 | 0.0126 |
| Equilibirium | No_Assortment | 2 | 519,530.2 | 195,097 | 145.197 | 1 | 0.0000 |
| No_Assortment | Sex_Invariant | 1 | 519,541.7 | 195,098 | 11.527 | 1 | 0.0007 |

**Supplementary Table 12: Model Fit Statistics (and Likelihood Ratio Tests) for EA (Non-cognitive)**

| Base | Comparison | *k* | -2LL | df | *Δ* 2LL | *Δ* df | *p* |
|---|---|---|---|---|---|---|---|
| Full | | 4 | 518,915.8 | 195,054 | | | |
| Full | Equilibirium | 3 | 518,923.0 | 195,055 | 7.193 | 1 | 0.0073 |
| Equilibirium | No_Assortment | 2 | 519,101.2 | 195,056 | 178.173 | 1 | 0.0000 |
| No_Assortment | Sex_Invariant | 1 | 519,102.2 | 195,057 | 1.039 | 1 | 0.3081 |

**Supplementary Table 13: Model Fit Statistics (and Likelihood Ratio Tests) for Depression**

| Base | Comparison | *k* | -2LL | df | *Δ* 2LL | *Δ* df | *p* |
|---|---|---|---|---|---|---|---|
| Full | | 4 | 522,445.5 | 195,171 | | | |
| Full | Equilibirium | 3 | 522,445.5 | 195,172 | 0.001 | 1 | 0.9809 |
| Equilibirium | No_Assortment | 2 | 522,448.8 | 195,173 | 3.279 | 1 | 0.0702 |
| No_Assortment | Sex_Invariant | 1 | 522,449.6 | 195,174 | 0.834 | 1 | 0.3610 |

**Supplementary Table 14: Model Fit Statistics (and Likelihood Ratio Tests) for Diabetes (Type 1)**

| Base | Comparison | *k* | -2LL | df | *Δ* 2LL | *Δ* df | *p* |
|---|---|---|---|---|---|---|---|
| Full | | 4 | 521,550.6 | 195,092 | | | |
| Full | Equilibirium | 3 | 521,551.9 | 195,093 | 1.232 | 1 | 0.2671 |
| Equilibirium | No_Assortment | 2 | 521,551.9 | 195,094 | 0.024 | 1 | 0.8773 |
| No_Assortment | Sex_Invariant | 1 | 521,551.9 | 195,095 | 0.007 | 1 | 0.9331 |

**Supplementary Table 15: Model Fit Statistics (and Likelihood Ratio Tests) for Diabetes (Type 2)**

| Base | Comparison | *k* | -2LL | df | *Δ* 2LL | *Δ* df | *p* |
|---|---|---|---|---|---|---|---|
| Full | | 4 | 521,940.1 | 195,005 | | | |
| Full | Equilibirium | 3 | 521,945.0 | 195,006 | 4.861 | 1 | 0.0275 |
| Equilibirium | No_Assortment | 2 | 521,945.0 | 195,007 | 0.001 | 1 | 0.9802 |
| No_Assortment | Sex_Invariant | 1 | 521,945.3 | 195,008 | 0.298 | 1 | 0.5850 |

**Supplementary Table 16: Model Fit Statistics (and Likelihood Ratio Tests) for Well-Being Spectrum**

| Base | Comparison | *k* | -2LL | df | *Δ* 2LL | *Δ* df | *p* |
|---|---|---|---|---|---|---|---|
| Full | | 4 | 522,074.8 | 195,109 | | | |
| Full | Equilibirium | 3 | 522,075.4 | 195,110 | 0.577 | 1 | 0.4474 |
| Equilibirium | No_Assortment | 2 | 522,075.6 | 195,111 | 0.206 | 1 | 0.6499 |
| No_Assortment | Sex_Invariant | 1 | 522,075.7 | 195,112 | 0.133 | 1 | 0.7157 |

Note: The p-values in the tables above have not been adjusted for multiple testing

### Supplementary Note 7 All polygenic index correlations

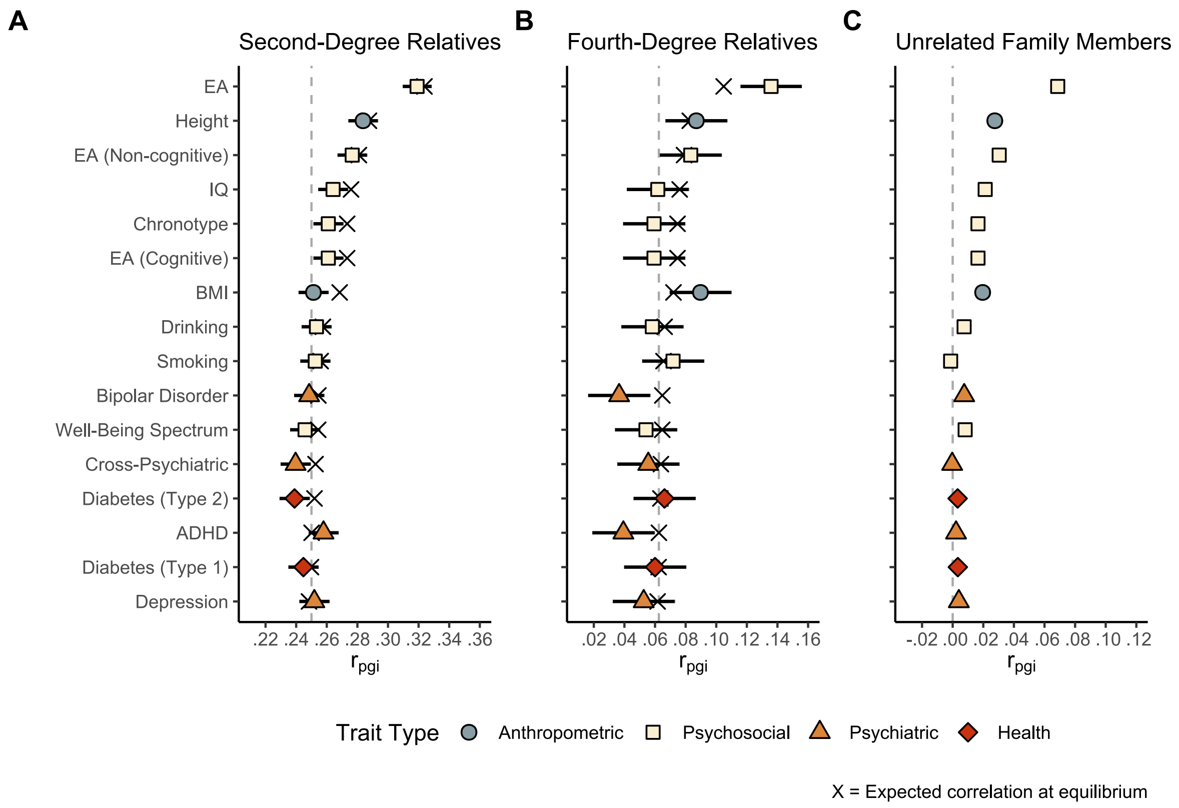

**Supplementary Figure 29** *Polygenic index correlations (with 95% Cis) between second-degree (N=35,923), fourth-degree (N=9,392), and unrelated family members (N=235,209).*

Supplementary Figure 29 shows the polygenic index correlations (95% CIs) among relatives that were not included in Fig. 3 in the main paper. Unrelated family members include all dyads in extended families that are not genetically related (i.e., relations that are mediated by a partner, such as in-laws, non-genetic uncles/aunts, partners, etc.).

The summary statistics and reproducible code for this figure is available at https://osf.io/dgw4r/.

Supplementary Table 17 lists correlations (and 95% confidence intervals) for all traits for all relations.

| Supplementary Table 17: Correlations (95% Confidence Intervals) | | | | | | | |
|---|---|---|---|---|---|---|---|
| Polygenic Index[a] | Partners | Parent-Offspring | Full Siblings | Second-Degree Relatives | Third-Degree Relatives | Fourth-Degree Relatives | Unrelated Family Members |
| EA (Total) | .138 | .560 | .558 | .319 | .197 | .136 | .069 |
| | *(.129, .147)* | *(.556, .564)* | *(.549, .567)* | *(.310, .328)* | *(.186, .208)* | *(.116, .156)* | *(.065, .073)* |
| Height | .072 | .533 | .531 | .284 | .151 | .087 | .028 |
| | *(.063, .081)* | *(.528, .537)* | *(.522, .540)* | *(.274, .293)* | *(.139, .162)* | *(.067, .107)* | *(.023, .032)* |
| EA (Non-cognitive) | .060 | .524 | .528 | .277 | .147 | .083 | .030 |
| | *(.051, .069)* | *(.520, .528)* | *(.518, .537)* | *(.267, .286)* | *(.135, .158)* | *(.063, .103)* | *(.026, .034)* |
| IQ | .050 | .521 | .513 | .264 | .140 | .062 | .021 |
| | *(.041, .059)* | *(.517, .525)* | *(.503, .522)* | *(.254, .274)* | *(.129, .152)* | *(.042, .082)* | *(.017, .025)* |
| Chronotype | .046 | .518 | .512 | .261 | .136 | .059 | .017 |
| | *(.037, .055)* | *(.514, .522)* | *(.502, .521)* | *(.251, .271)* | *(.125, .148)* | *(.039, .079)* | *(.012, .021)* |
| EA (Cognitive) | .046 | .518 | .512 | .261 | .136 | .059 | .017 |
| | *(.037, .055)* | *(.514, .522)* | *(.502, .521)* | *(.251, .271)* | *(.125, .148)* | *(.039, .079)* | *(.012, .021)* |
| BMI | .036 | .513 | .500 | .251 | .140 | .090 | .020 |
| | *(.027, .045)* | *(.509, .517)* | *(.490, .509)* | *(.242, .261)* | *(.128, .151)* | *(.070, .110)* | *(.016, .024)* |
| Drinking | .015 | .506 | .515 | .253 | .134 | .058 | .007 |
| | *(.006, .024)* | *(.501, .510)* | *(.506, .525)* | *(.244, .263)* | *(.123, .146)* | *(.038, .078)* | *(.003, .012)* |
| Smoking | .012 | .509 | .514 | .252 | .127 | .072 | -.001 |
| | *(.003, .021)* | *(.505, .513)* | *(.504, .523)* | *(.243, .262)* | *(.115, .138)* | *(.052, .092)* | *(-.005, .003)* |
| Bipolar Disorder | .009 | .504 | .505 | .248 | .129 | .037 | .008 |
| | *(-.000, .018)* | *(.500, .508)* | *(.495, .515)* | *(.239, .258)* | *(.118, .141)* | *(.016, .057)* | *(.003, .012)* |
| Well-Being | .009 | .500 | .511 | .246 | .126 | .054 | .008 |
| Spectrum | *(-.000, .018)* | *(.495, .504)* | *(.502, .521)* | *(.236, .256)* | *(.114, .137)* | *(.034, .074)* | *(.004, .012)* |
| Cross-Psychiatric | .005 | .497 | .497 | .240 | .112 | .056 | -.000 |
| | *(-.004, .014)* | *(.493, .502)* | *(.487, .507)* | *(.230, .249)* | *(.100, .123)* | *(.035, .076)* | *(-.004, .004)* |
| Diabetes (Type 2) | .004 | .497 | .495 | .239 | .115 | .066 | .003 |
| | *(-.005, .013)* | *(.493, .501)* | *(.485, .505)* | *(.229, .249)* | *(.104, .127)* | *(.046, .086)* | *(-.001, .007)* |
| ADHD | .000 | .500 | .503 | .258 | .118 | .039 | .002 |
| | *(-.009, .009)* | *(.496, .505)* | *(.494, .513)* | *(.248, .267)* | *(.107, .130)* | *(.019, .059)* | *(-.002, .006)* |
| Diabetes (Type 1) | -.001 | .499 | .499 | .245 | .121 | .060 | .003 |
| | *(-.010, .008)* | *(.494, .503)* | *(.489, .509)* | *(.235, .254)* | *(.110, .133)* | *(.040, .080)* | *(-.001, .007)* |
| Depression | -.003 | .498 | .512 | .252 | .123 | .053 | .004 |
| | *(-.012, .006)* | *(.494, .503)* | *(.502, .521)* | *(.242, .262)* | *(.112, .135)* | *(.032, .073)* | *(-.000, .008)* |

[a] *Sorted by correlation between partners*

#### Supplementary Note 8 Information about the polygenic indices

**Supplementary Table 18: Polygenic indices**

| Polygenic Index | GWAS sample size | Overlapping SNPs |
|---|---|---|
| Educational Attainment (EA)[22] | 765283 | 908380 |
| Non-Cognitive EA[25] | 510795 | 879419 |
| Cognitive EA[25] | 257700 | 879419 |
| Intelligence[26] | 269867 | 911938 |
| Well-Being Spectrum[27] | 2311180 | 800700 |
| Height[28] | 4080687 | 1041747 |
| Body Mass Index (BMI)[29] | 695648 | 802341 |
| Cigarettes per day (Smoking)[30] | 245876 | 907384 |
| Drinks per week (Drinking)[30] | 941280 | 402099 |
| Type 1 Diabetes[31] | 520580 | 911412 |
| Type 2 Diabetes[32] | 659316 | 827710 |
| Cross-Psychiatric Disorders[33] | 734126 | 688620 |
| Bipolar Disorder[34] | 413466 | 912724 |
| Broad Depression[35] | 500199 | 256913 |
| ADHD[36] | 225534 | 902783 |
| Chronotype[37] | 697828 | 879419 |

Overlapping SNPs refers to the number of single nucleotide polymorphisms (SNP) that were available in both the GWAS summary statistics and in the genotype data in MoBa. It is therefore the number of SNPs used to calculate the polygenic indices. Non-cognitive EA and cognitive EA are based on GWAS-by-subtraction as described in [25]. In short, the polygenic index for educational attainment is split into that related to intelligence and that unrelated to intelligence.
